# N-terminal phosphorylation of Huntingtin: A molecular switch for regulating Htt aggregation, helical conformation, internalization and nuclear targeting

**DOI:** 10.1101/358234

**Authors:** Sean M. DeGuire, Francesco. S. Ruggeri, Mohamed-Bilal Fares, Anass Chiki, Urszula Cendrowska, Giovanni Dietler, Hilal A. Lashuel

**Author notes:** These authors contributed equally to this work (Francesco. S. Ruggeri, Mohamed-Bilal Fares).

## Abstract

Phosphorylation of exon1 of the Huntingtin protein (Httex1) has been shown to play important roles in regulating the structure, toxicity and cellular properties of N-terminal fragments and the full-length Huntingtin protein. Here, we investigated and compared the effect of bona fide phosphorylation at S13 and/or S16 on the structure, aggregation, membrane binding, and subcellular properties of mutant Httex1-Q18A. We show that serine phosphorylation at either S13 or S16 strongly disrupts the amphipathic α-helix of the N-terminus, inhibits the aggregation of mutant Httex1 and prompts the internalization and nuclear targeting of Httex1 preformed aggregates. In synthetic peptides phosphorylation at S13 and/or S16 strongly disrupted the amphipathic α-helix of the N-terminal 17 residues (Nt17) of Httex1 and Nt17 membrane binding. Our studies on peptides bearing a different combination of phosphorylation sites within Nt17 revealed a novel phosphorylation-dependent switch for regulating the structure of Httex1 involving crosstalk between phosphorylation at T3 and S13 or S16. Together, our results provide novel insights into the role of phosphorylation in regulating Httex1 structure and function in health and disease and underscore the critical importance of identifying enzymes responsible for regulating Htt phosphorylation and their potential as therapeutic targets for the treatment of Huntington’s disease.

## Introduction

Huntington’s disease (HD) is a fatal neurodegenerative disorder that results from a CAG repeat expansion (>36) in the first exon of the *IT-15* gene encoding the Huntingtin protein (Htt) (1). Although the length of the CAG-encoded polyQ tract seems to directly correlate with disease severity and Htt protein aggregation propensity (2–4), other sequence features including post-translational modifications (PTMs) have been shown to influence the aggregation, cellular properties and toxicity of Htt and therefore may contribute to HD pathogenesis (5–11). The identification of PTMs that promote or protect against Htt-induced toxicity could lead to the development of new drugs for the treatment of HD by modulating these modifications and their downstream effects.

N-terminal fragments of Htt have been found in nuclear inclusion bodies in HD patients (12), and overexpression of exon1 of Htt (Httex1) with expanded polyQ repeats alone is sufficient to recapitulate many characteristics of HD in mice (13), suggesting that N-terminal Htt fragments may play a critical role in the pathogenesis of HD (14,15). These observations combined with the fact that these fragments represent more tractable systems for structure-function studies have made N-terminal Htt fragments the focus of many studies aimed at elucidating the structural basis of Htt aggregation and toxicity.

Httex1 is composed of 3 domains: the Nt17 domain comprising the first 17 amino acids, the polyQ domain and the proline-rich domain, all of which play important roles in the misfolding, oligomerization and/or fibrillization of Httex1 and full-length Htt (4,16–22). *In vitro* biophysical studies suggest that Httex1 aggregation, although polyQ dependent, occurs through a two-step mechanism that involves an association of the Nt17 domain to form homo-oligomeric species characterized by increased α-helical content which then undergoes conformational changes to form β-sheet-rich fibrils that grow through monomer addition (23). Other studies suggest that the Nt17 interaction with the polyQ domain influences the structure of Httex1 in ways that favor the population of aggregation-prone conformational intermediates (17,24,25). Consistent with these studies, N-terminal Htt fragments lacking Nt17 show reduced aggregation rates compared with those containing Nt17 with similar polyQ repeat lengths (19,26). In addition to its role in Htt aggregation, this domain has been implicated in the regulation of full-length protein-protein interactions (21,27–30), subcellular localization (10), and toxicity (31).

The Nt17 domain comprises several residues that have been shown to undergo phosphorylation (T3 (24), S13, and S16 (6,32)) or other post-translational modifications such as acetylation (5) ubiquitination and SUMOylation (33) (K6, K9, and K15) in HD brains, transgenic animals and in cell culture models of HD. Increasing evidence suggests that these PTMs influence Htt structure, aggregation (23,31), subcellular localization (32,34), toxicity (5), and protein-protein interactions (6), and may represent important molecular switches for regulating these properties in health and disease (35). However, dissecting the specific effects of single or multiple modifications has been challenging, because the main enzymes involved in regulating many of these modifications remain unknown. For example, serine phosphorylation of Httex1 has been reported to be mediated by IκB kinase (IKK) (6) and casein kinase 2 (CK2) (32), however, the efficiency and selectivity of these enzymes are too poor to make them reliable tools for efficient manipulation of phosphorylation at these residues *in vivo* or for the generation of homogeneously phosphorylated recombinant Htt proteins *in vitro*. Because of these challenges, most studies investigating the effects of phosphorylation to date have relied on the overexpression of phosphomimetic (S→D/E) or phosphoresistant (S→A) mutant forms of Htt. In cell culture models, the introduction of phosphomimetic mutations at both S13 and S16 has been consistently shown to modify the levels of Httex1 ubiquitination and acetylation, increase Htt nuclear localization(32), reduce its toxicity and influence its clearance (6). However, overexpression of these mutants in animal models of HD has led to conflicting results. Whereas in *Drosophila*, the overexpression of (S13D/S16D) 120Q mutant Htt (mHtt) enhanced toxicity compared to unmodified mHtt (36,37), whereas the introduction of the S13D/S16D mutations in full-length Htt with expanded polyQ repeats (97Q) in a BACHD transgenic mouse model reversed mutant-Htt induced pathology and disease phenotype. In terms of motor deficits, psychiatric-like deficits, and aggregate formation, S→D mutant BACHD mice were indistinguishable from non-transgenic mice (31). Moreover, recent studies showed overexpression of single phosphomimetic mutations of S13 and S16 in human cells (using a bimolecular fluorescence complementation assay) or in Drosophila was sufficient to inhibit or abolish the aggregation of Httex1 (38), and inhibitions of PP1 phosphatase which modulates Nt17 phosphorylation was shown to decrease both the aggregation and neurotoxicity (38). However, even in the same study, the effect of the phosphomimetics varied depending on the expression system used with phosphomimicking mutations at S13 and S16 leading to decreased or increased aggregation, depending on whether they are expressed in the larval eye imaginal discs or Drosophila or in adult dopaminergic neurons. The discrepancy between these studies could be due to the fact that the use of mutations to mimic or abolish phosphorylation does not allow for assessing the dynamics of phosphorylation and the role of potential cross-talk between the phosphorylation at T3, S13 and S16. Furthermore, increasing evidence shows that the phosphomimicking mutations do not reproduce all aspects of protein phosphorylation (24,39–41). We recently showed that the T3D mutation did not fully reproduce the inhibitory effect of T3 phosphorylation on the aggregation of Httex1 (24).

To address these limitations, we took advantage of recent advances made by our group that enable the site-specific introduction of post-translational modifications within residues the Nt17 domain in the context of a tag-free native wild-type (22Q) (42) and mutant (42Q) Httex1 using protein semisynthesis (24). We used these strategies to produce, in milligram quantities, single and double-phosphorylated (pS13, pS16 and pS13/pS16) forms of Httex1 with polyQ repeats both above and below the pathogenic threshold of HD. With these proteins in hand, we performed systematic studies to address the following questions: 1) What is the effect of phosphorylation at each or both serine residues on the aggregation and fibril morphology of Httex1? 2) Is the effect of serine phosphorylation-dependent on the polyQ repeats length? 3) Does the introduction of phosphomimetic mutations at S13 and/or S16 reproduce the effects of phosphorylation at these residues on Httex1 secondary structure and aggregation? and, 4) How does phosphorylation at S13 and/or S16 affect the internalization and subcellular targeting of Httex1 aggregates in primary neurons?

## Results

### Semi-synthesis of Httex1 protein

Expressed protein ligation requires a nucleophilic cysteine residue at the N-terminus of the C-terminal ligation fragment and a thioester at the C-terminus of the N-terminal fragment. Httex1 does not contain cysteine residues. In such cases, a transient alanine to cysteine mutation is usually introduced to create a ligation site and the native alanine residue and native protein sequence can be restored by reductive desulfurization of the ligated product. However, no alanine residues exist between the desired phosphorylation sites (S13 and S16) and the polyQ repeat domain. Therefore, we created a ligation site by substituting the first glutamine residue of the polyQ domain with a cysteine (Q18C) to generate wild-type (22Q) and mutant (42Q) Httex1. After desulfurization, this leads to the introduction of a Q→A mutation at residue 18 (Q18A) in all proteins generated using this semisynthetic strategy. In the absence of kinases that allow for site-specific phosphorylation at S13 and/or S16, this is the only way to introduce site-specific phosphorylation with minimal perturbation of the native sequence Figure 1A. Therefore, as a control, Httex1 proteins without serine modifications, but bearing this mutation (Q18A-Httex1, **Unmodified Httex1-Q_22/42_**) were prepared using the same semisynthetic strategy outlined above and were always used for direct comparison with the modified proteins in all our biophysical studies. To make sure that the presence of the Q18A mutation does not affect the aggregation kinetics of mutant Httex1, we compared the aggregation of **Httex1-42Q Q18A** to the aggregation of **Httex1-43Q** at different concentrations (5, 10 and 20 μM). Figure 2A shows that the aggregation kinetics as determined by monitoring the loss of Httex1 monomers are not affected by the Q18A mutation. In addition, examination of the fibrils formed by **Httex1-43Q** and **Httex1-42Q Q18A** revealed virtually identical fibril morphologies for both proteins (Figure 2B).

**Figure 1:**
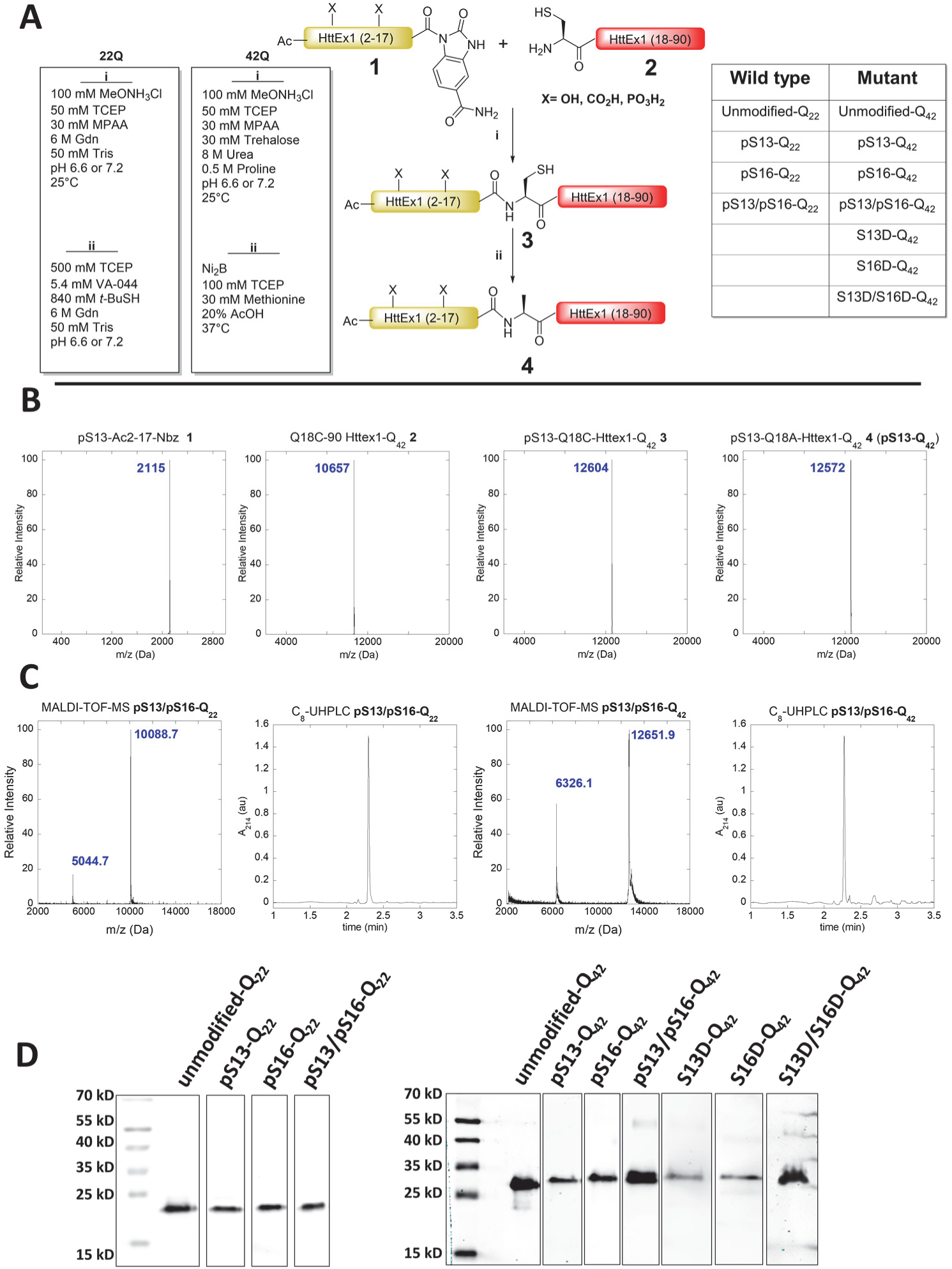
Semi-synthesis of Httex1 phosphorylated at S13 and/or S16. (**A**) A schematic depiction of the semi-synthetic strategy used to produce phosphorylated (pS) wild-type (22Q) and mutant (42Q) Q18A Httex1 and the corresponding phosphmimicking mutants. (**B**) The ESI-MS analyses of starting materials, intermediates and products of native chemical ligation and desulfurization for pS13-Q_42_ are shown. **(C)** Final purity characterization of Httex1 pS13/pS16-Q_22_ and Httex1 pS13/pS16-Q_42_ using MALDI-TOF mass spectrometry and UHPLC **(D)** Western blot analysis characterization of the proteins produced using this semisynthetic strategy and listed in (**A)**.

**Figure 2:**
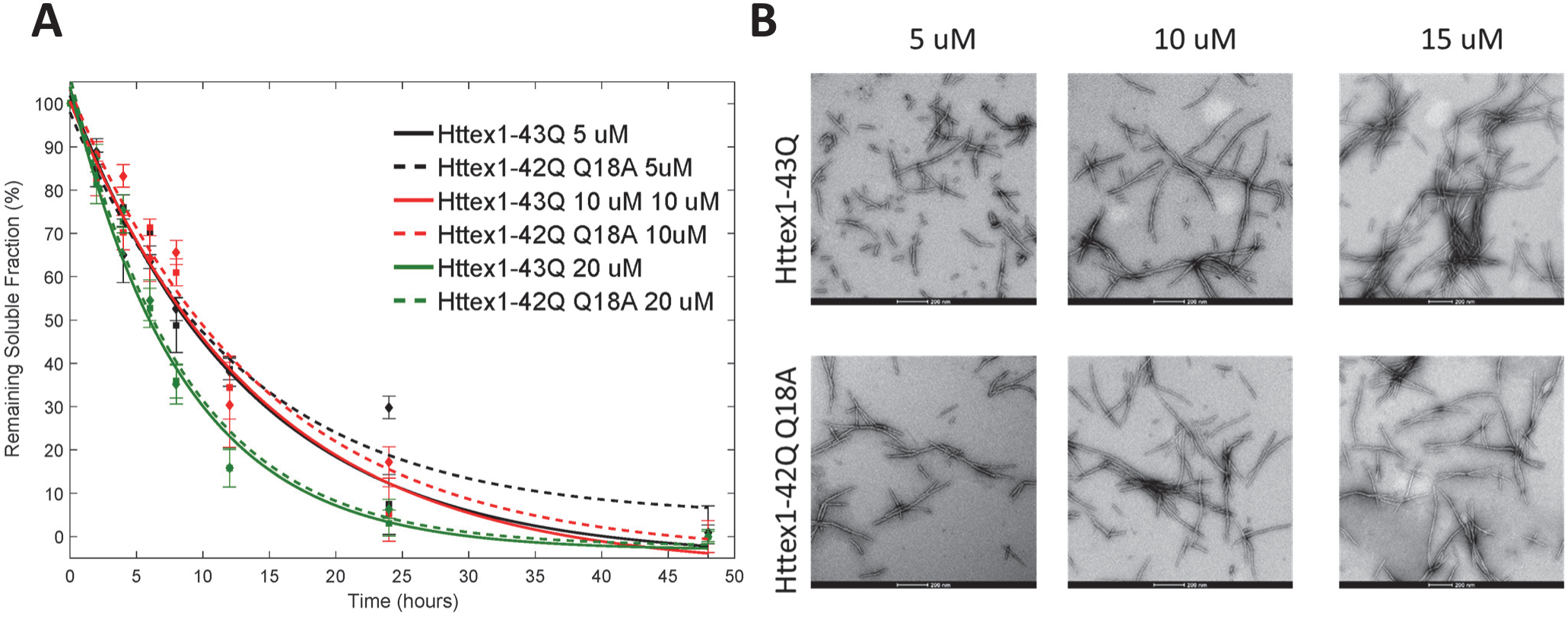
The substitution of Q18 by alanine (Q18A) to facilitate the semisynthesis of phosphorylated Httex1 proteins via NCL does not alter the kinetics of aggregation or morphology of the fibrils formed by mutant Httex1. (**A**) Comparison of the aggregation kinetics of Httex1-43Q and Httex1-42Q bearing the Q→A mutation at residue 18 (Q18A) at 5 μM, 10 μM and 20 μM as determined by the UHPLC sedimentation assay. **(B)** TEM analysis of Httex1-43Q and Httex1-42Q (Q18A) at different concentrations (scale bars are 200 nm).

To elucidate the role of phosphorylation at these residues, we generated Httex1 phosphorylated at single serine residues (pS13 or pS16) and both residues (pS13/pS16). To determine whether S-D substitutions would reproduce the effect of phosphorylation at these sites, we also prepared proteins with the corresponding phosphomimetic mutations S13D, S16D, and S13D/S16D using the same semisynthetic strategy. Figure 1A shows a schematic depiction of our semisynthetic strategy and a list of the semisynthetic Httex1 proteins generated and used in this study.

To produce the proteins required for this study, a series of seven N-terminal peptides were prepared as N-acyl-benzimidazolinones (NBz) that incorporated the desired phosphorylated residue(s) (43). Previous studies have shown that Httex1 undergoes methionine removal and subsequent A2 acetylation *in vivo* (5). Consequently, all Nt17 peptides used in this study were N-terminally acetylated. To prepare wild-type (22Q) Httex1 proteins, the Q18C 18–90 22Q Httex1 fragment was expressed in *E. coli* and subsequently purified using an intein-based expression strategy that was recently developed in our laboratory for the production of Httex1 (24). N-terminal thiazolidine products formed from the undesired condensation of cellular electrophiles were removed by hydrolysis at low pH and at 37°C in the presence of excess methoxyamine (44). Native chemical ligation reactions of this fragment (**2**) with each peptide (**1**) were conducted under reducing and denaturing conditions at neutral pH in the presence of 4-mercaptophenylacetic acid (MPAA) and yielded the desired ligation products at room temperature (24,42). Unexpectedly, under these conditions, the NBz group of peptides containing S16 phosphorylation was found to convert rapidly to side-products via hydrolysis and aminolysis before significant product formation was observed. For ligations with pS16 and pS13/pS16 peptides, the omission of methoxyamine treatment prevented aminolysis and the slow addition of neat peptide-NBz (2.0 EQ over 4 hours) maintained sufficient amounts of active thioester for NCL to proceed rapidly. Each ligated protein was then treated with radical reductive desulfurization conditions (45). The resulting Q18A Httex1 proteins were purified by RP-HPLC and their purity was established by SDS-PAGE, Reverse Phase Ultra High-performance liquid chromatography (RP-UHPLC), LC-MS and MALDI-TOF-MS.

Similarly, mutant Httex1 proteins were prepared by native chemical ligation of the same peptides with recombinant Q18C 18–90 Httex1 (**2**) containing an expanded polyQ repeat (42Q). These ligations were more challenging because the unreacted recombinant C-terminal fragment (**2**) and the ligation products (**3**) exhibited increased tendency to aggregate during the ligation and desulfurization reactions before completion of the desired transformations. To overcome these challenges, we utilized ligation conditions that we have recently developed a method based on the use of a combination of strongly denaturing conditions with additives that are known to inhibit protein and polyQ peptide aggregation (24). Under these conditions, the ligation reactions proceeded to completion before the aggregation of the proteins could occur. Nickel Boride mediated reductive desulfurization was found to be compatible with an acidic water/organic solvent system that keeps Httex1 protein soluble even at 37°C, the temperature required for efficient reduction (46). The purity and chemical identity of the mutant proteins after RP-HPLC purification was established by SDS-PAGE, RP-UHPLC, ESI-MS and MALDI-TOF-MS (Figure 1).

### Single Phosphorylation at S13 or S16 slows the aggregation of mutant Httex1

To evaluate whether single phosphorylation or phosphomimetic mutations at S13 or S16 inhibit the aggregation of mutant (42Q) Httex1, we compared the aggregation properties of **unmodified Httex1-Q_42_, pS13 Httex1-Q_42_, pS16 Httex1-Q_42_, S13D Httex1-Q_42_**, and **S16D Httex1-Q_42_** Httex1 proteins. The lyophilized proteins were first treated with a previously established disaggregation protocol to allow for solvation in aqueous buffer and to disrupt any preformed aggregates (47). The proteins were then dissolved in PBS, the pH was adjusted to 7.3 and any remaining preformed aggregates were removed by filtration (100 kDa cutoff filter, Merck Millipore). The concentration of each filtrate was adjusted to 2.5 µM using a polyQ repeat length-specific calibration curve based on the UHPLC profiles of samples of known concentration as determined by amino acid analysis (48). The aggregation of the proteins was assessed using a sedimentation assay and monitoring the loss of soluble protein. Sample aliquots were periodically collected over the course of several days from solutions incubated at 37°C. Following the removal of Httex1 aggregates from these aliquots via sedimentation, the remaining fraction of soluble protein was measured by quantitative UHPLC analysis of the supernatant relative to the initial protein concentration.

Complete aggregation and conversion of monomeric **unmodified Httex1-Q_42_** into insoluble aggregates occurred within 24 hours (Figure 3A). At this time point, all phosphorylated mutant Httex1 proteins showed reduced aggregation compared to the unmodified Httex1, with **pS13 Httex1-Q_42_** and **pS16 Httex1-Q_42_** proteins being approximately 40% aggregated and only 80% of the proteins converted into insoluble aggregates after 48 hours. To obtain an estimate of the rate of aggregation (loss of monomer), the data obtained for each protein aggregation assay were fit to an exponential decay function (48–50). The effect of the single phosphorylation is confirmed when the approximated relative aggregation rates were calculated from the sedimentation data, phosphorylation of either serine (for **pS13 Httex1-Q_42_** and **pS16 Httex1-Q_42_**) resulted in aggregation rates only 45% of the rate of **unmodified-Q_42_** (Figure 3B). This inhibitory effect was not reproduced by the S → D substitutions at these residues. The Httex1 phosphomimetics **S13D Httex1-Q_42_** and **S16D Httex1-Q_42_** exhibited similar aggregation rates and degrees of aggregation as the **unmodified Httex1-Q_42_** (Figure 3A/B).

**Figure 3:**
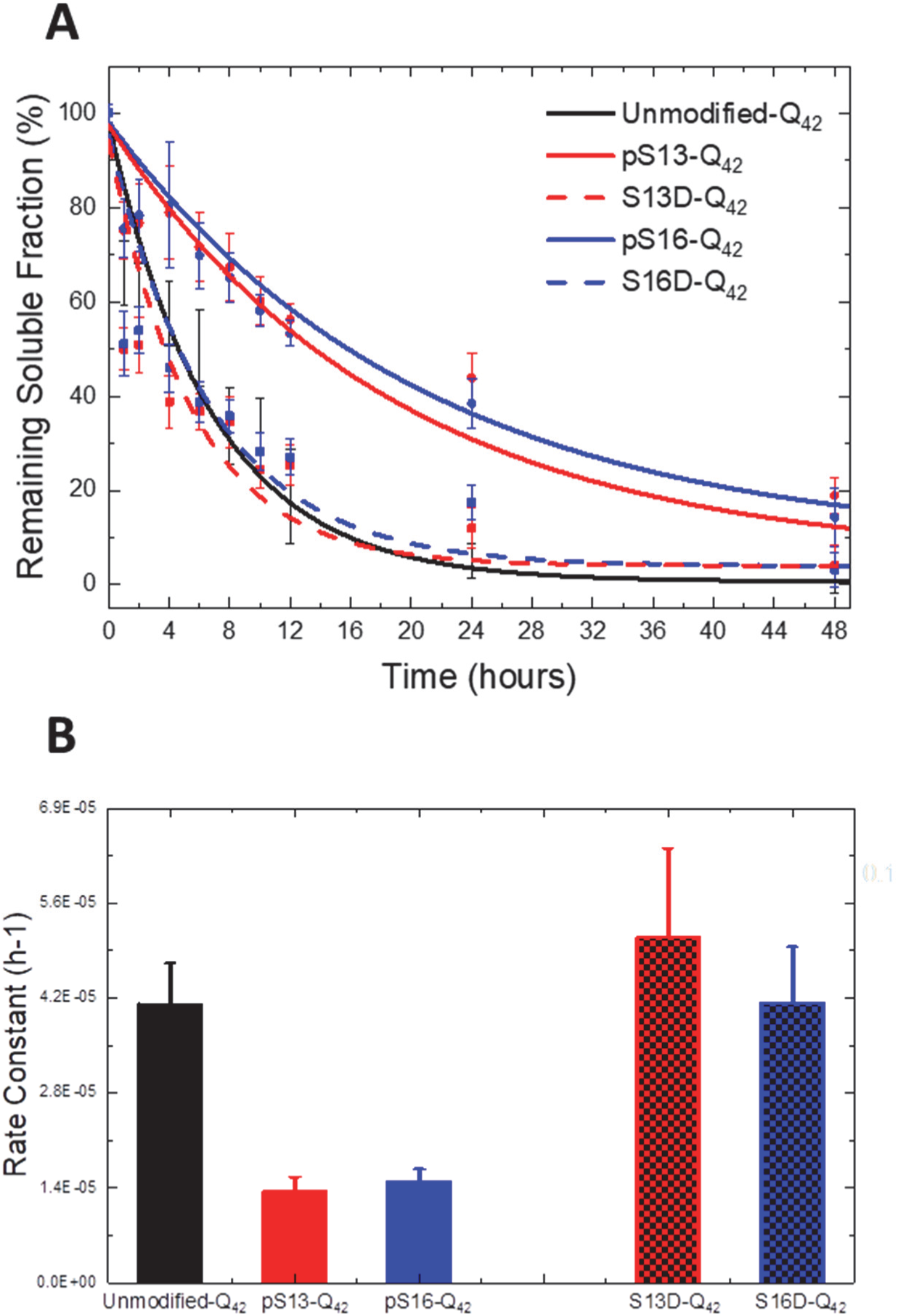
Phosphorylation at S13 or S16, but not pseudophosphorylation, results in a significant retardation of mutant Httex1 aggregation. **(A)** Aggregation analysis of single phosphorylated and phosphomemitic mutant (42Q) Httex1 Q18A (pS13, pS16, S13D and S16D) at 2.5 µM using the UHPLC sedimentation assay and **(B)** calculated aggregation rates.

By CD spectroscopy, the **unmodified Httex1-Q_42_** protein underwent a shift in secondary structure that was consistent with a transition from unstructured (λ_min_ 205 nm) to predominantly β-sheet structure (λ_min_ 212–215 nm) within 24 hours (Figure 4). For proteins bearing single phosphomimetic mutations (**S13D Httex1-Q_42_** and **S16D Httex1-Q_42_**), the CD spectra at 24 hours were indistinguishable from those of the **unmodified Httex1-Q_42_** (Figure 4B), consistent with the nearly complete loss of soluble protein by sedimentation analysis. In contrast, CD analysis of proteins bearing a single phosphorylation (**pS13 Httex1-Q_42_** and **pS16 Httex1-Q_42_**) at this time point showed only a decrease in the CD signal without a significant shift in λ_min_.

**Figure 4:**
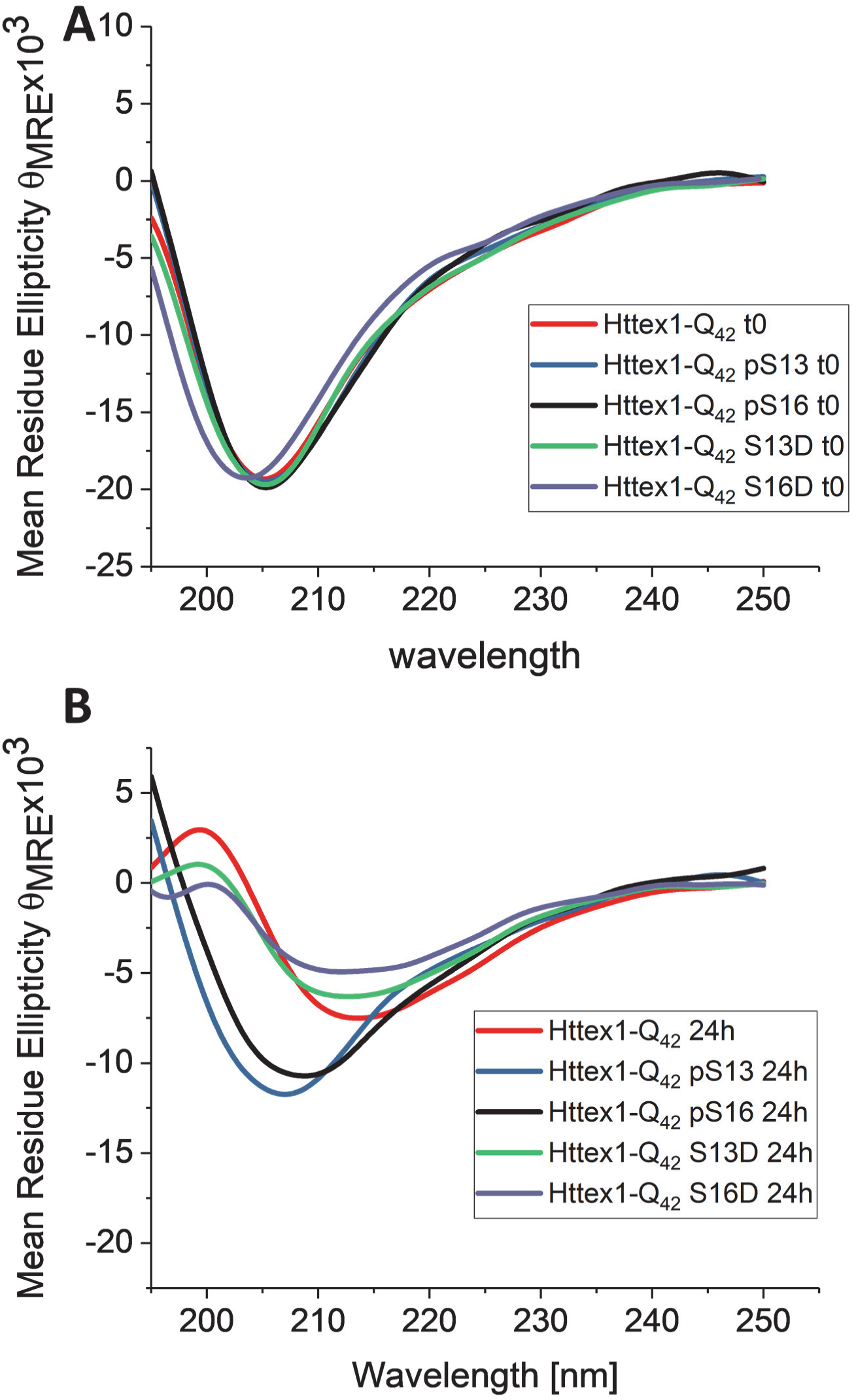
Only unmodified Httex1 and the S13D and S16D mutants are able to undergo a transition from random coil to β-sheet structure upon aggregation. Circular dichroism spectra of singly phosphorylated (at S13 or S16) mutant Httex1 and the corresponding phosphomemitic proteins at 0h **(A)** and 24h **(B).**

To determine the effect of phosphorylation on the morphological properties of mutant Httex1 fibrils, we performed AFM high-resolution imaging and statistical analysis on the aggregates formed by each protein. Consistent with the *in vitro* aggregation data, we observed that the phosphorylated proteins **pS13-Q_42_** and **pS16-Q_42_** formed significantly fewer aggregates compared with the **unmodified-Q_42_**. However, the data presented in Figure 5 focuses on the quantitative comparison of the fibrils morphology, as determined by assessing changes in the height and length distribution of the aggregates formed by each Httex1 variant, which can be qualitatively related to differences in the extent of aggregation of the different proteins. The AFM images of the **unmodified Httex1-Q_42_** as early as five hours of incubation revealed mature fibrillar aggregates with average heights of 6.3±0.5 nm and average lengths of 176±83 nm (Figure 5). The phosphorylated proteins, **pS13 and pS16-Httex1-Q_42_**, produced significantly smaller aggregates with average heights of 4.7±0.6 nm and 5.1±0.6 nm and average lengths of 67±20 nm and 70±22 nm, respectively. Only the aggregate length, but not the height of these structures was observed to increase after 48 hours. Interestingly, mutant Httex1 protein bearing a single phosphomimetic mutation (**S13D Httex1-Q_42_** and **S16D Httex1-Q_42_**) formed aggregates at five hours with average heights of 5.8±0.6 nm and 5.2±0.8 nm and average lengths of 77±29 nm 82±38 nm, respectively (Figure 5B/C). After 48 hours, the height of the S→D mutant Httex1 aggregates had increased notably to sizes comparable to **unmodified Httex1-Q_42,_** suggesting that the introduction of phosphomimetic mutation at a single serine residue only slightly influence the final morphology of the fibrils, consistent with the ability of the **WT, S13D, and S16D Httex1-Q_42_** to form β-sheet rich aggregates (Figure 4) (51). In contrast, phosphorylation at S13 or S16 resulted in fibrils with significantly reduced heights compared to the **unmodified Httex1-Q_42_**. Interestingly, **Httex1-Q_42_** bearing *bona fide* phosphorylation or phosphomimetic mutations at S13 or S16 exhibited similar fibril length distributions after 48 hours. Together, these findings combined with the CD data suggest that phosphorylation at S13 or S16 residues significantly reduce the rate of fibril formation and inhibits transition to β-sheet rich conformation, but does not alter the overall length distribution of the Httex1 fibrils. In contrast, single S→D substitution at these residues does not inhibit the aggregation of **Httex1-Q_42_** or alter the ability of the protein to form β-sheet rich aggregates.

**Figure 5:**
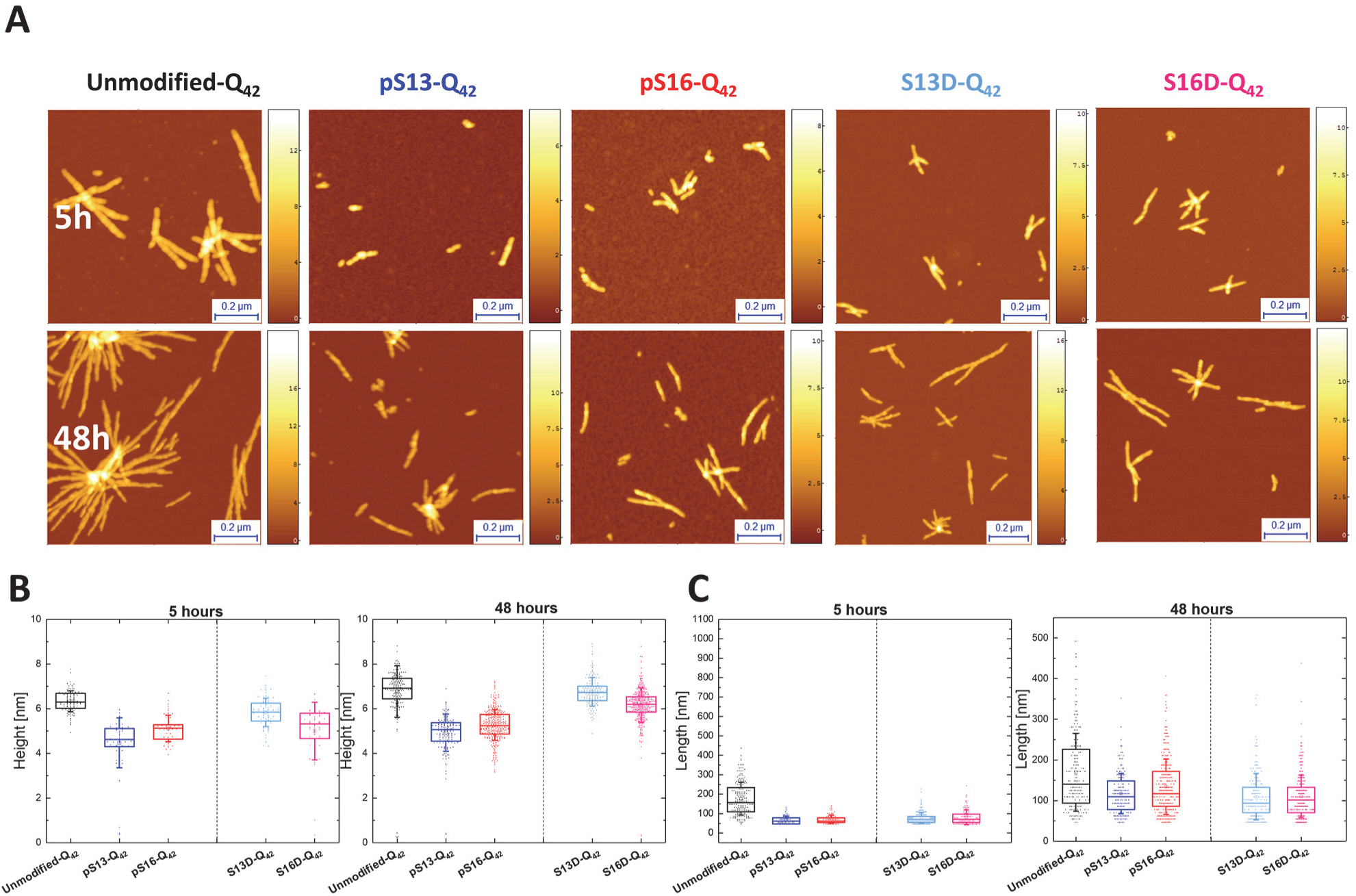
Phosphorylation at S13 or S16 results in the formation of fibrils with significantly reduced heights compared to the unmodified Httex1-Q42 and the corresponding phosphomemitic mutants. (**A**) AFM images of aggregates formed by mutant 42Q Httex1 Q18A phosphorylated at S13 or S16 and the corresponding phosphomimetic mutants. Quantitative analysis of the height (**B**) and length (**C**) of these aggregates measured from AFM images. (Z scale bars are in nm). All scatter plots of height differ with a statistical significance of at least p < 0.05. All scatter plots of the length of the modified aggregates differ from the unmodified with a statistical significance of at least p < 0.05.

### Diphosphorylation at S13/S16 inhibits mutant Httex1 aggregation

Phosphorylation at both S13/S16 was shown to play important roles in Htt pathology in animal models of HD (31). Different studies using phosphomimetic mutations (S13D/S16D) or Httex1-like peptides showed that phosphorylation at both residues (pS13/pS16) influence the aggregation of Httex1 and reverses Htt’s pathology in an HD mouse model (23,31,32). However, chemical and enzymatic tools were not available to determine the structural consequences of the S13/S16 phosphorylation on full length and untagged Httex1 aggregation. Therefore, we sought to use our semisynthetic strategy to produce mutant Httex1 bearing *bona fide* phosphorylation of both of these residues (pS13/pS16) and compare its aggregation properties to the commonly used corresponding phosphomimetic mutant (S13D/S16D).

As shown in Figure 6A, phosphorylation of both S13 and S16, **pS13/pS16 Httex1-Q_42,_** resulted in a dramatic inhibition of Httex1-43Q aggregation as determined by both the sedimentation assay and TEM (Figure 6A and C). The corresponding phosphomimetic mutant (**S13D/S16D Httex1-Q_42_**) exhibited significantly slower aggregation compared to the unmodified protein or the single phosphomimetic mutants (S13D or S16D, Figure 3), which individually did not have any effect on mutant Httex1 aggregation. After 48 hours of incubation, 82% of the **pS13/pS16 Httex1-Q_42_** protein remained soluble compared to 60% for **S13D/S16D Httex1-Q_42_ (Figure 6A)**, These observations were confirmed by CD spectroscopy, which revealed no significant reduction or shift in CD spectra of **pS13/pS16 Httex1-Q_42_** after 48 hours **(Figure 6B),** suggesting that the protein exists predominantly in monomeric and/or non-structured low molecular weight oligomeric forms. In the case of the disphosphomimetic protein (**S13D/S16D Httex1-Q_42_**) mutant, a significant reduction in the CD signal was observed **(Figure 6B)**, consistent with the loss of soluble protein due to the increased aggregation propensity of this protein and detection of fibrils by TEM **(Figure 6C)**. These results demonstrate that phosphorylation at both serine residues (**pS13/pS16 Httex1-Q_42_)** results in a strong inhibitory effect on mutant Httex1 aggregation and that the phosphomimetic mutations at both of these residues also result in significant inhibition of mutant Httex1 aggregation, but do not fully reproduce the effect of *bona fide* phosphorylation at these residues.

**Figure 6:**
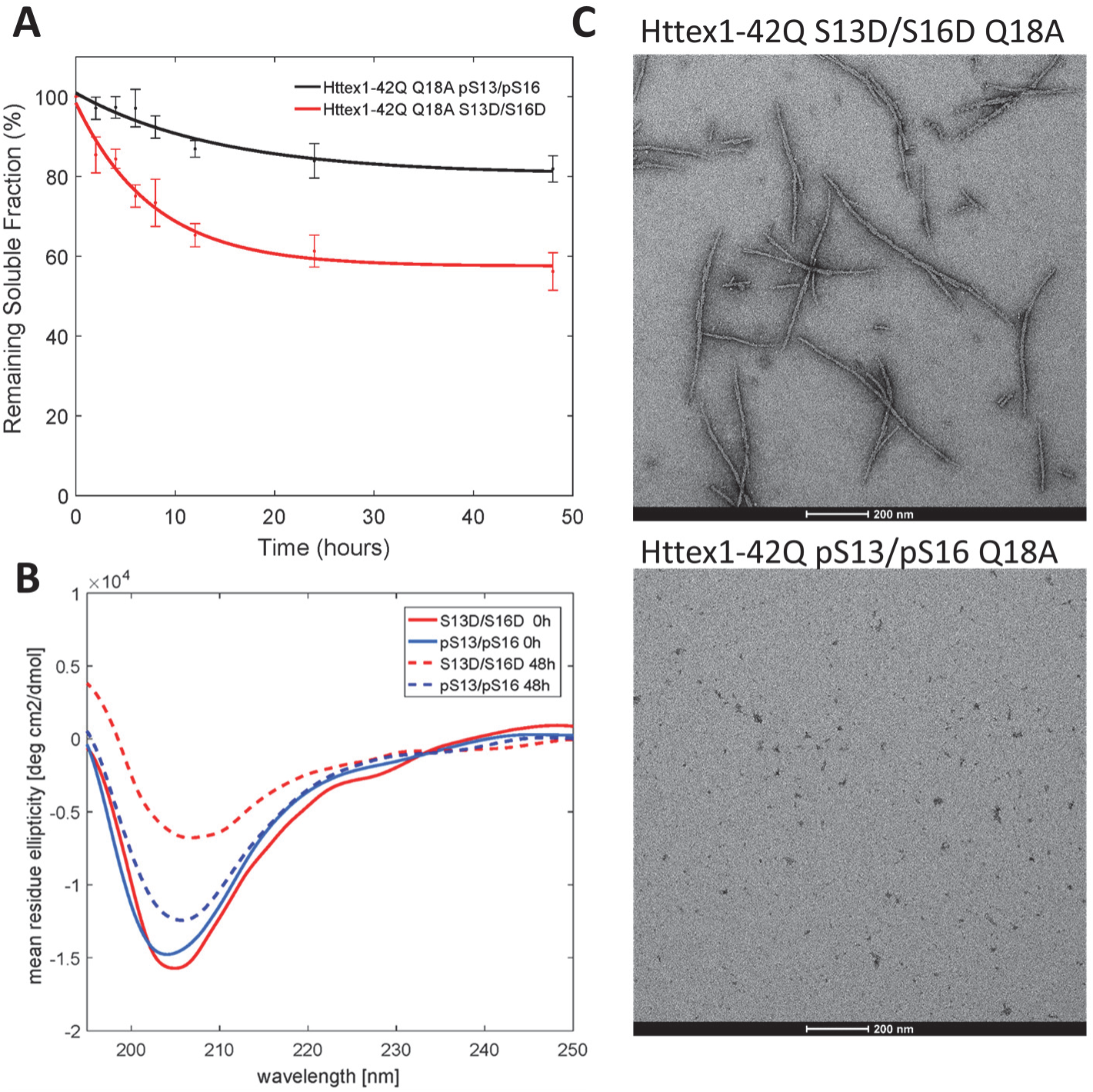
Phosphorylation at both S13 and S16 inhibits the aggregation of Httex1-Q42 Q18A, whereas the S13D/S16D Httex1-Q42 Q18A exhibits a marked reduction in aggregation but retains the ability to form fibrils. **(A)** Comparative analysis of the aggregation of diphosphorylated Q18A (pS13/pS16) and diphosphomemitic (S13D/S16D) mutant (42Q) Httex1 at 2.5 µM using the UHPLC sedimentation assay and **(B)** CD analysis of the samples at 0 and after 48 hours of incubation at room temperature. **(C)** TEM images of the final aggregation time point (48 hours). Scale bars are 200 nm.

### Serine phosphorylation at S13 and/or S16 inhibits wild-type Httex1 aggregation

Previous reports from our group have shown that Httex1 proteins with polyQ repeat lengths well below the pathological threshold (23Q) are able to aggregate and form oligomers and fibrils *in vitro*, albeit at a much lower rate, over a long period of time (days) and to a lesser extent compared with mutant forms of Httex1 with polyQ repeats of >36Q (26,42). To evaluate the effect of serine phosphorylation on the biophysical properties of wild-type Httex1, the aggregation of **unmodified Httex1-Q_22_, pS13 Httex1-Q_22_, pS16 Httex1-Q_22_** and **pS13/pS16 Httex1-Q_22_** Httex1 was monitored using the *in vitro* sedimentation assay described above (26). **Figure 7A** shows that serine phosphorylation at S13 or S16 reduced the rate and extent of wild-type Httex1 aggregation. For **unmodified Httex1-Q_22_** Httex1, 75% of the protein aggregated after four weeks of incubation at 37°C. In contrast, in the case of **pS13 Httex1-Q_22_** or **pS16 Httex1-Q_22_** Httex1, approximately 40% of the protein was aggregated, indicating that phosphorylation at either residue reduced the extent of Httex1 aggregation to a similar degree. A much stronger effect was observed when both S13 and S16 residues were phosphorylated (**pS13/pS16 Httex1-Q_22_**), where < 20% of the protein was aggregated after four weeks. The amount of soluble protein (monomers and non-sedimentable oligomers) remaining at each intermediate time point was always greater for the phosphorylated Httex1 proteins indicating a reduction of the aggregation rate as well. As in the case of **Q_42_** proteins (*vide supra*), the approximated relative aggregation rates were calculated from the sedimentation data. The rates (ks^−1^) were found to be approximately 50% for **pS13 Httex1-Q_22_** and **pS16 Httex1-Q_22_** Httex1 and approximately 20% for **pS13/pS16 Httex1-Q_22_** Httex1 (**Figure 7B**) relative to the rate of **unmodified Httex1-Q_22_** Httex1 protein. The inhibitory effect of serine phosphorylation on wild-type Httex1 aggregation appeared to be additive, with the diphosphorylated Httex1 exhibiting approximately twice the effect of single phosphorylation at S13 or S16.

**Figure 7:**
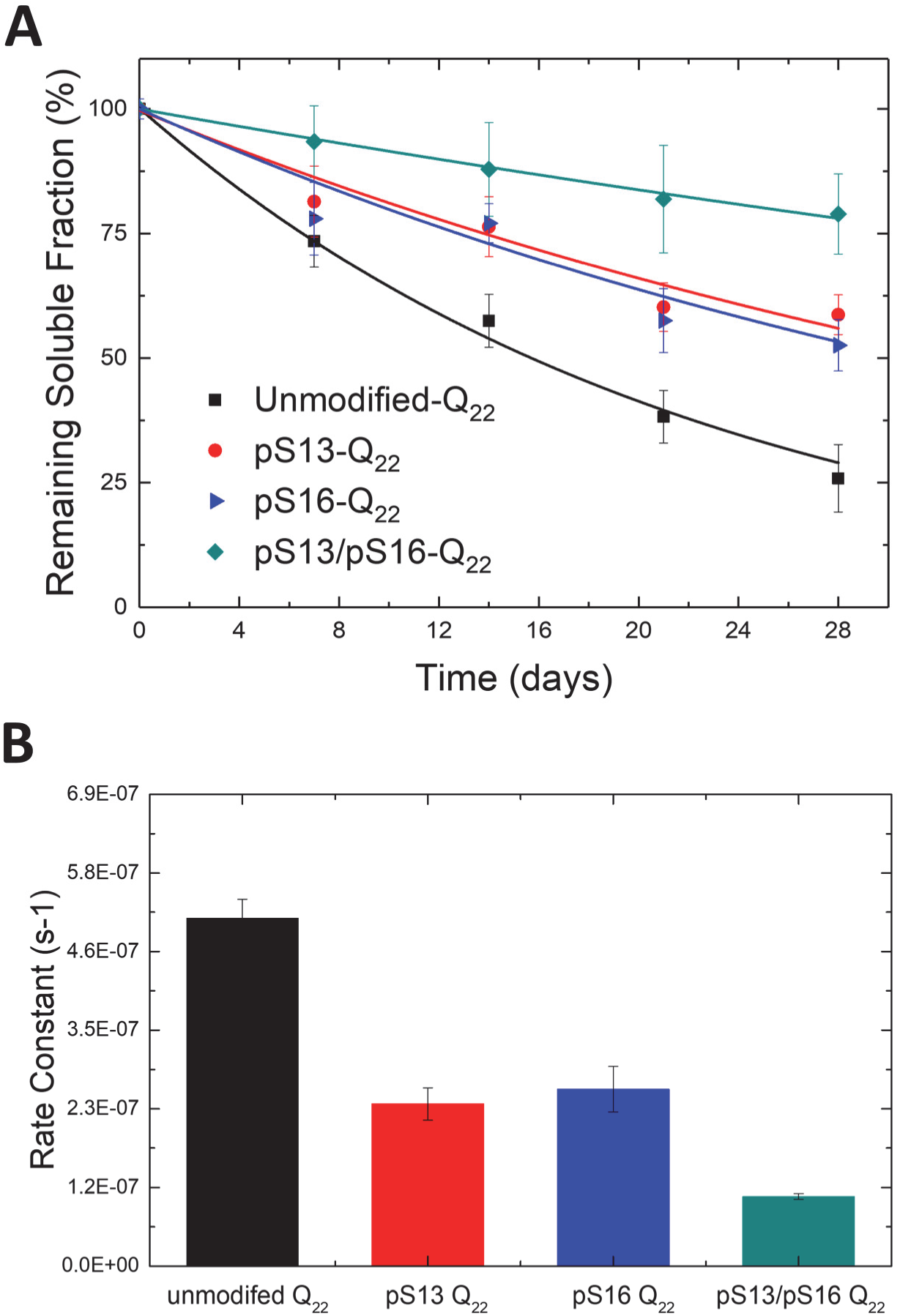
Serine phosphorylation at S13 and/or S16 inhibits wild-type Httex1 aggregation. **(A)** Comparison of the aggregation of unmodified, singly phosphorylated and diphosphoryated WT (22Q) Q18A Httex1 protein at 5 µM as determined using the UHPLC sedimentation assay **(B)** calculated aggregation rates.

### Serine phosphorylation disrupts the amphipathic α-helix of the Htt Nt17 domain

To determine the effect of phosphorylation on the conformational properties of Nt17, we assessed the effect of phosphorylation at S13 and/or S16 on the helix propensity of Nt17 in buffered solutions or upon binding to membranes. The transient helical nature of Nt17 combined with the fact that the vast majority of the Httex1 sequence beyond Nt17 exists in a disordered conformation makes any significant secondary structural changes within Nt17 of the full-length Httex1 challenging to monitor by CD spectroscopy. Therefore, we used synthetic N-terminal peptides bearing the first 19 amino acids of Httex1 (MATLEKLMKAFESLKSFQQ), and where S13, S16 or both residues were phosphorylated.

The Nt17 exists predominantly in a disordered conformation but exhibits a high propensity to form transient helical conformations. This is consistent with their CD spectra, which show a minimum at 200–205 nm and small shoulder at 218–225 nm (**Figure 8A).** Importantly, serine phosphorylation at S13 and/or S16 resulted in a decrease in α-helical content as indicated by the reduction of this shoulder minimum in comparison to the unmodified Nt19 peptide **(Figure 8A**). These findings are consistent with previous reports using Httex1-like model peptides containing the Nt17 peptide (32). Recently, we showed that threonine 3 (T3) phosphorylation results in significant stabilization of the α-helical conformation of Nt17 and a significant increase in Nt17 α-helical contents measured by CD spectroscopy and NMR (24). To determine if phosphorylation at S13 and/or S16 could disrupt the stable helical conformation of Nt17 induced by phosphorylation at T3, we synthesized peptides that are phosphorylated at T3 and S13, T3 and S16 and T3 and both S13 and S16 and assessed their conformation by CD. Phosphorylation at S13 resulted in a stronger destabilization of the pT3 induced α-helix formation compared to phosphorylation at S16 (**Figure 8B**). Phosphorylation of all the threonine and serine residues (T3, S13 and S16) resulted in complete disruption of the Nt19 helical conformation. Indeed, the triphosphorylated peptide showed a CD spectrum that exhibits even less α-helical content than the unmodified peptide, indicating that phosphorylation at both S13 and S16 completely reverses the effect of T3 phosphorylation on the secondary structure of Nt17 (**Figure 8B**). Taken together, these results reveal a novel phosphorylation-dependent conformational switch whereby the α-helical conformation of Nt17 is regulated by phosphorylation/dephosphorylation of T3 or by crosstalk between T3 and S13 and/or S16 phosphorylation.

**Figure 8:**
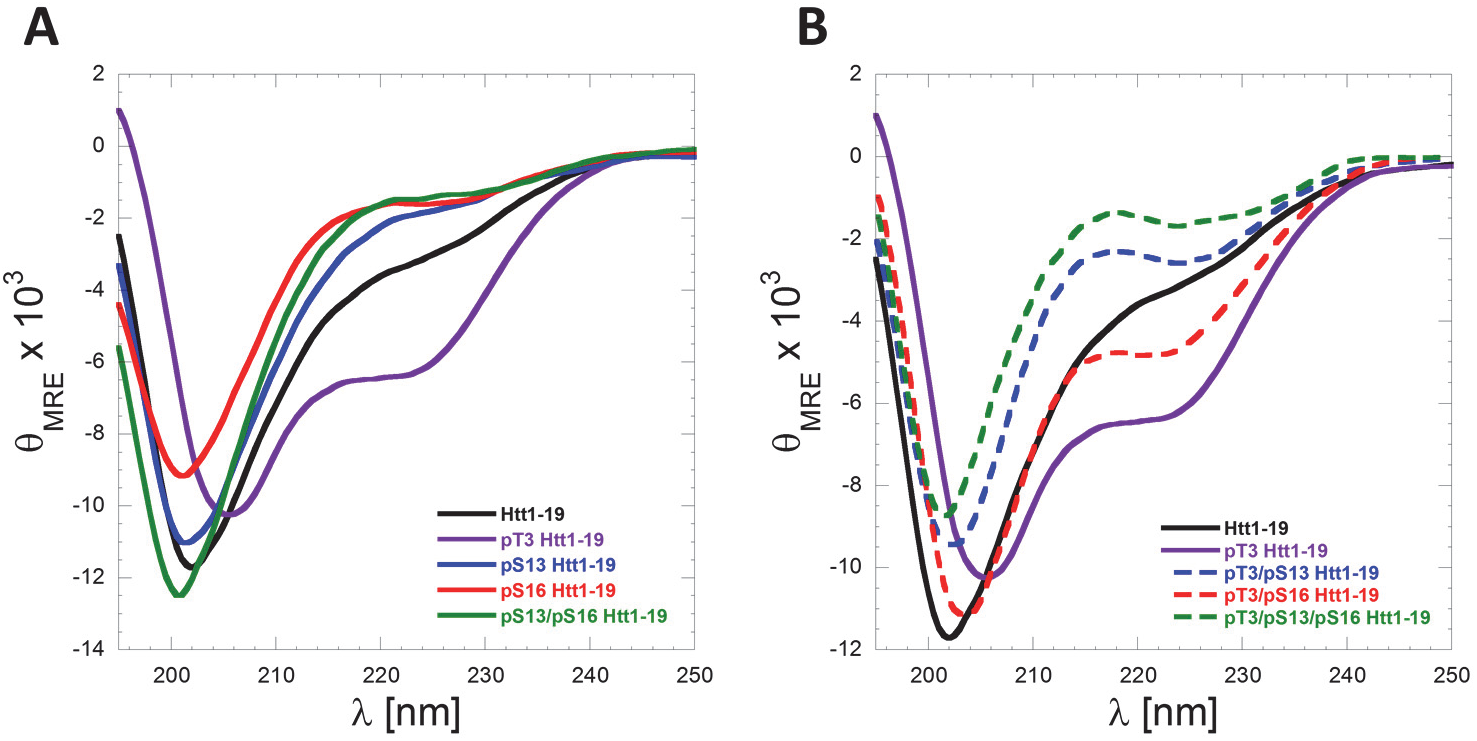
Serine phosphorylation at S13 and/or S16 disrupts the amphipathic α-helix of Nt17 and reverses pT3-induced α-helix formation of Nt17. **(A)** Far-UV CD spectra (θ_MRE=_ Mean Residue Ellipticity) of Htt1-19 peptides bearing single (pT3, pS13 or pS16) or multiple phosphorylated (pS13/pS16) residues at 60 µM (pH 7.4). **(A)** Phosphorylation of T3 increases the helicity of Nt17, whereas phosphorylation at S13, S16 or both residues reduces the helical propensity of Nt17. **(B)** Phosphorylation at S13, S16, or both residues disrupts pT3-induced α-helix formation of Nt17. To allow for better visualization and comparison of the data and the effect of phosphorylation at S13 and/or S16 on pT3-induced **α-helix** formation, the Far-UV-CD spectra of Htt 1–19 and Htt 1–19 pT3 were re-plotted again in (B).

The Nt17 has been shown to adopt α-helical structure upon membrane binding (52) and act as an anchor for Httex1 to membranes and mediate its aggregation on membranes (53). To determine the effect of phosphorylation on Nt17 membrane binding properties, we assessed the effect of the phosphorylation at S13 and/or S16 and their corresponding phosphomimetic variants on the binding of the Nt17 peptide to phosphatidylglycerol (POPG) as a membrane model. Large POPG unilamellar vesicles (LUVs) with an average diameter of 100 nm were used at 5 molar equivalent. As expected, the unmodified Nt17 peptide formed an α-helix upon incubation with LUVs **(Figure 9A)**. Phosphorylation at S13 or S16 significantly inhibited the random coil to α-helix transition by Nt17 in the presence of lipid vesicles with phosphorylation at S13 inducing the strongest effect. When both S13 and S16 residues are phosphorylated (pS13/pS16), the Nt17 peptide failed to adopt α-helical conformations in the presence of membranes and exhibited CD spectra that are similar to that of Nt17 in the absence of membranes. **(Figure 9A/C)**. In contrast, the introduction of the phosphomimetic mutation (S→D) at S13 or S16 or both residues did not influence the ability of Nt17 to adopt an α-helical conformation in the presence of POPG LUVs **(Figure 9B/C)**. These results demonstrate that the phosphomimetics do not reproduce the effect of phosphorylation on Nt17 conformation upon membrane binding, highlighting the importance of using *bona fide* phosphorylation to study the effect of this modification on the structural properties of Httex1.

**Figure 9:**
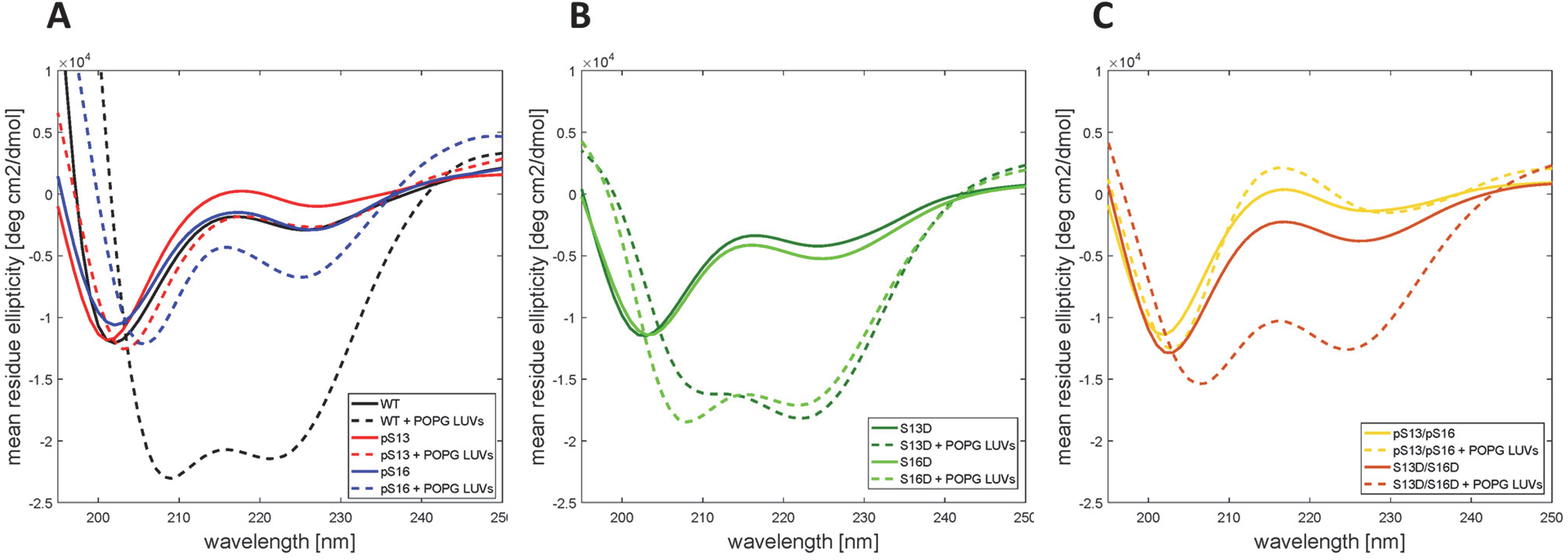
Phosphorylation at S13 and/or S16, but not the phosphomimetic (S13D and/or S16D) mutations, inhibits helix formation and binding of the Nt17 peptides to Large POPG unilamellar vesicles (LUVs) **(A)** Far-UV CD spectra of WT, pS13 and pS16 Nt17 in the presence and absence of LUVs. **(B)** Far-UV CD spectra of S13D and S16D in the presence and absence of LUVs. (C) Far-UV CD spectra of the diphosphorylated (pS13/pS16) Nt17 and the corresponding phosphomimetic mutant(S13D/S16D) in the presence and absence of LUVs.

### Serine phosphorylation prompts the internalization and subcellular targeting of Httex1 in neurons

Increasing lines of evidence from cellular models (54–57), animal models (58–60) (70–72), and recently from HD patients (61) support a role for the cell to cell transmission and propagation of Htt aggregates in the pathogenesis of HD. Therefore, we sought to investigate the effect of serine phosphorylation on Httex1 aggregate internalization and subcellular targeting in striatal primary neurons. To achieve this goal, we first generated preformed fibrillar species (PFFs) of each of the different proteins by incubating monomers at a high concentration (60–100 uM) at 37°C for several days. The preparations were then briefly sonicated to generate small fibrillar fragments, and then characterized by TEM. Dot blot analysis was also performed to establish that the relative concentrations of the different preparations were similar. Next, we treated primary striatal neurons with 0.5 µM PFFs (Httex1-Q_22_ or Httex1-Q_42_) with single/double phosphoserine residues, or with the unmodified PFFs as a control. Notably, immunocytochemistry using the MW8 anti-Httex1 antibody (C-term) showed that whereas unmodified PFFs were mostly distributed outside the cellular membrane with very rare instances of internalization of few foci of PFFs, neurons treated with phosphoserine-bearing Httex1-Q_22_ or Httex1-Q_42_ PFFs displayed, in addition to these distributions, uptake of large quantities of PFFs that appeared to fill up the cytoplasm in a homogenous manner (Figure 10A, 11A). Orthogonal projections of high-resolution confocal z-stacks verified the intracellular localization of the phosphoserine aggregates in treated neurons, in contrast to the unmodified PFFs, which mostly appeared to sit on the extracellular membrane (**Figure 10B, 11B**).

**Figure 10:**
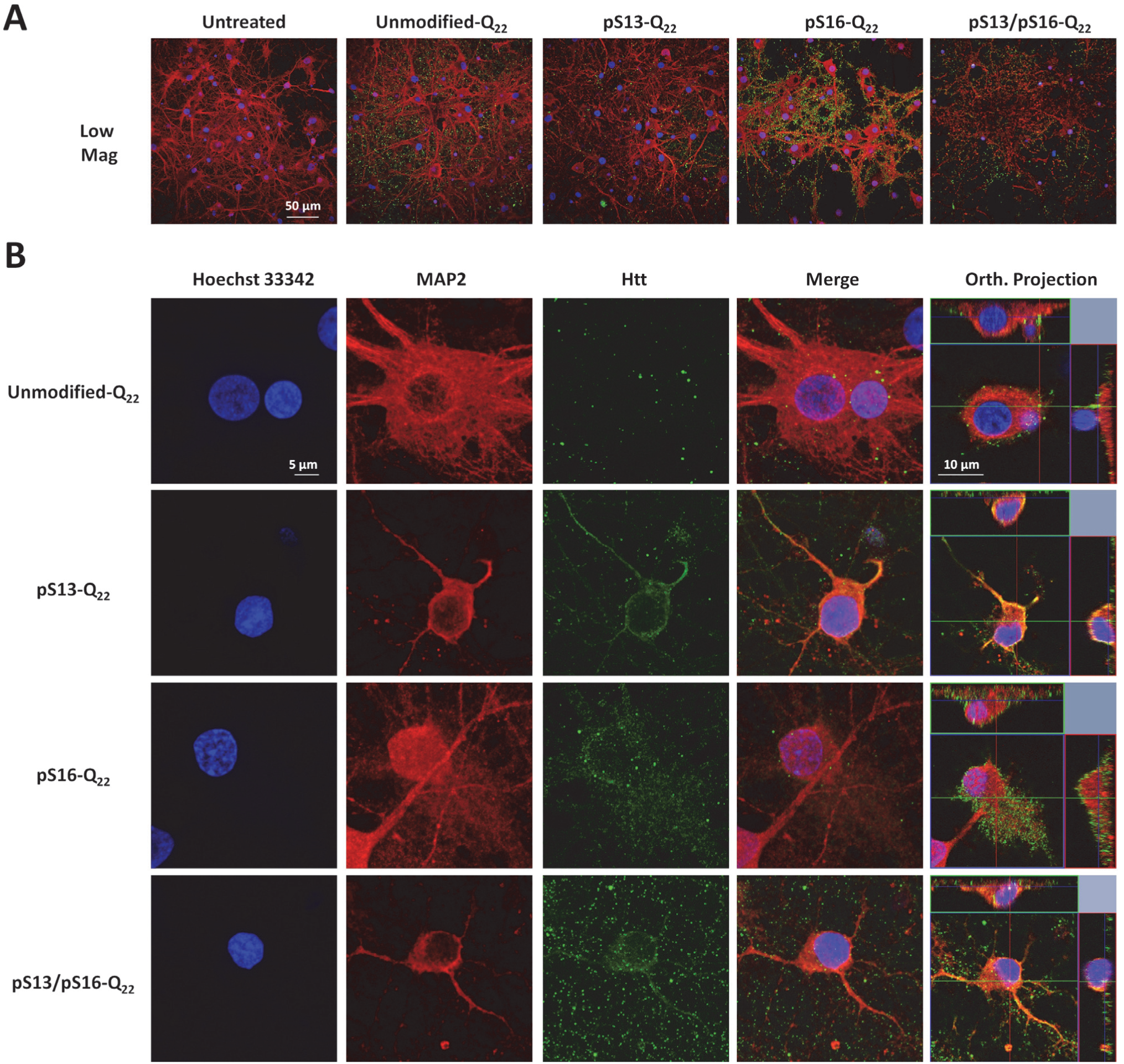
Effect of phosphorylation at S13 and/or S16 on the internalization of wild-type (22Q) Httex1 PFFs in striatal neurons. Primary rat striatal neurons treated with 0.5 µM of unmodified or PTM Q_22_ PFFs were immuno-stained using the MW8 anti-total Htt antibody. Staining for MAP2 and Hoechst-33342 was also performed to reveal neuronal cell bodies and nuclei, respectively. (**A**) Low magnification imaging showing that the MW8 antibody equivalently detected unmodified and PTM Q_22_ Httex1 PFFs, in contrast to the untreated condition. (**B**) High magnification confocal imaging of individual neurons showing that while **umodified-Q22** PFFs are mostly not internalized, some neurons treated with **pS13-Q22, pS16-Q22** and **pS13/pS16-Q22** Httex1 PFFs exhibit prominent internalization of PFFs, as established by orthogonal projection of z-stacks (Orth. Projection).

**Figure 11:**
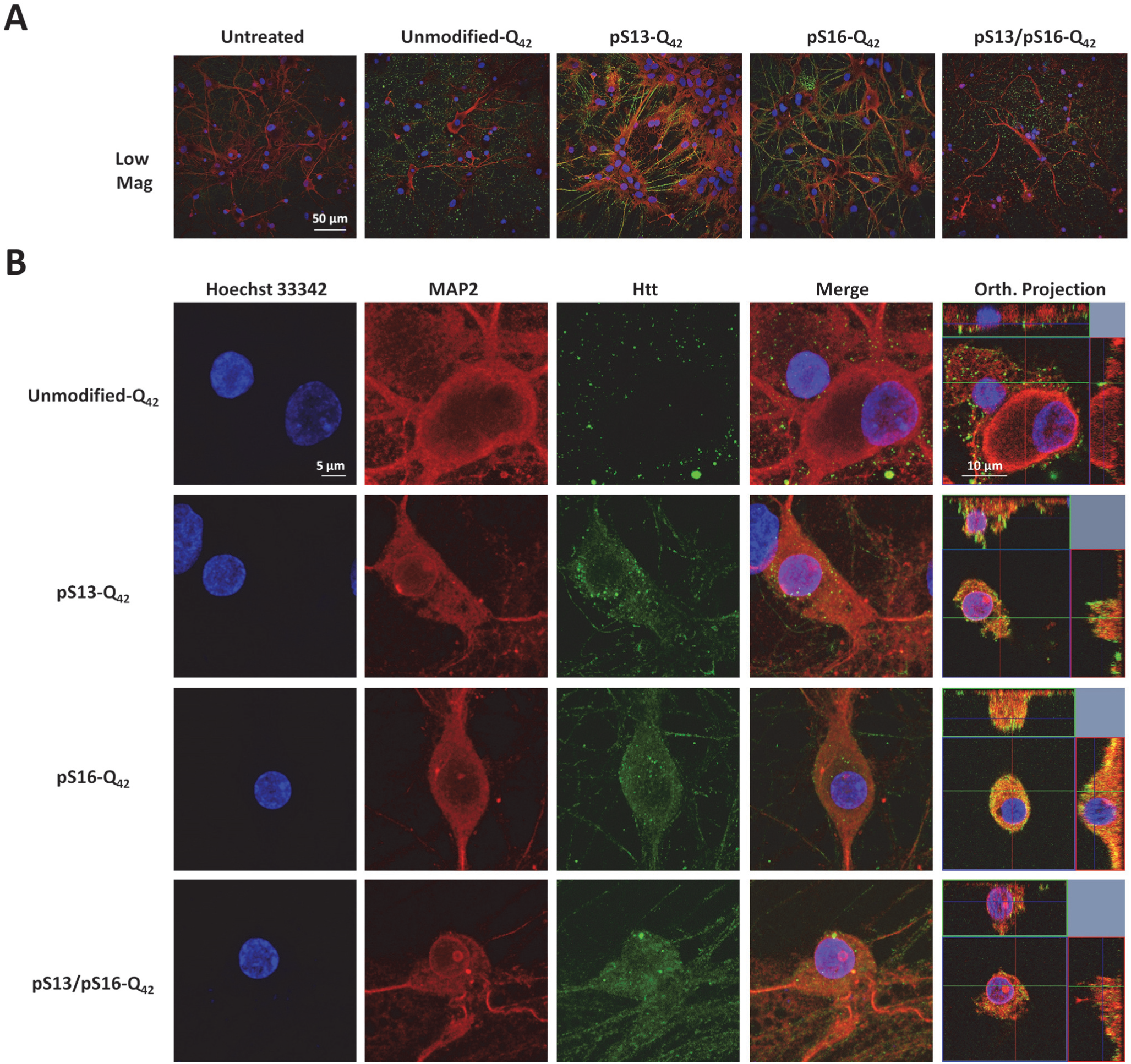
Effect of phosphorylation at S13 and/or S16 on the internalization of mutant (Q42) Httex1 PFFs in striatal neurons. Primary rat striatal neurons treated with 0.5 μM of unmodified or PTM Q42 PFFs were immuno-stained using the MW8 anti-total Htt antibody. MAP2 and Hoechst-33342 staining were also performed to reveal neuronal cell bodies and nuclei, respectively. (**A**) Low magnification imaging showing that the MW8 antibody equivalently detected unmodified and PTM Q_42_ PFFs in contrast to the untreated condition. (**B**) High magnification confocal imaging of individual neurons showing that while **umodified-Q42** PFFs are mostly not internalized, some neurons treated with **pS13-Q42, pS16-Q42** and **pS13/pS16-Q42** Httex1PFFs exhibit prominent internalization of PFFs, as established by orthogonal projection of z-stacks (Orth. Projection).

In addition, some of the neurons that had internalized PFFs bearing serine phosphorylation also exhibited additional subcellular targeting to the nuclear compartment (**Figure 12A**). To dissect the effect of single versus double phosphorylation on the nuclear targeting of Httex1 aggregates, we quantified the number of neurons exhibiting nuclear Httex1 localization upon treatment with unmodified versus the different phosphoserine bearing PFF variants. Notably, we found that both single and double phosphorylation of Nt17 significantly enhanced the targeting Httex1-Q_22_ and Httex1-Q_42_ PFFs to the nuclear compartment compared with unmodified PFFs (**Figure 12B**), with neurons treated with pS13/pS16 double phosphorylated PFFs exhibiting a slight tendency toward increased nuclear localization. Nevertheless, no statistically significant differences were noted among the three phosphorylated variants, indicating that a single phosphorylation event at either S13 or S16 is sufficient to prompt the nuclear enrichment of Httex1-Q_22_ or Httex1-Q_42_ aggregates.

**Figure 12:**
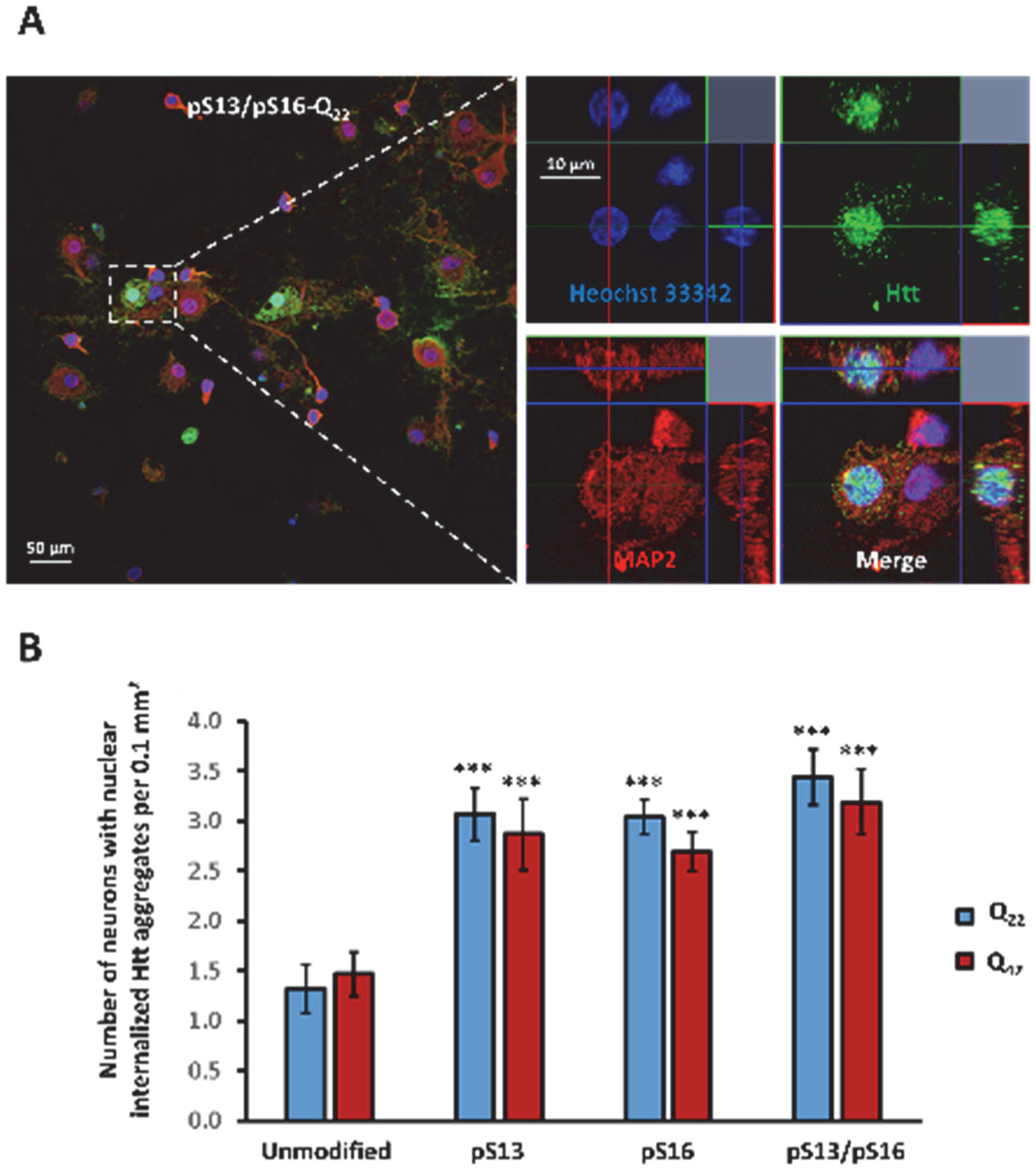
Effect of phosphorylation at S13 and/or S16 on the subcellular localization of mutant (Q42) PFFs in striatal neurons. (**A**) Immunostaining using the MW8 antibody revealing nuclear localization of internalized **pS13/pS16-Q22** PFFs in a proportion of neurons. A low magnification view is shown in the left panel, and the right panel shows high-resolution images of an individual neuron exhibiting nuclear localization. (**B**) Significantly more neurons treated with phosphorylated Q_22_ and Q_42_ Httex1 PFFs exhibit nuclear localization per 0.1 mm2 compared with the controls treated with unmodified Q_22_ or Q_42_ Httex1 PFFs. The asterisks denote P<0.001 determined by one-way ANOVA and Scheffé post hoc statistical analysis.

## Discussion

Protein phosphorylation is a dynamic PTM that allows precise temporal control of protein structure, function and subcellular localization. Phosphorylation of proteins implicated in neurodegenerative diseases such as Tau in Alzheimer’s disease, α-synuclein in Parkinson’s disease and TDP-43 in Amyotrophic lateral sclerosis seems to correlate with the formation of pathological aggregates and disease progression (62). In Huntington’s disease, phosphorylation of N-terminal serine and threonine residues has been detected in human brains, transgenic models and cell culture models of HD (5,31,63,64). However, the role of phosphorylation in regulating the structure, aggregation and HD pathology remains poorly understood, primarily due to the lack of tools that enable site-specific and efficient modulation of phosphorylation *in vitro* and *in vivo*.

The observation that phosphomimetic mutations at serine 13 and 16 attenuate disease progression and accumulation of Htt aggregates in the BACHD mouse model highlights the potential role of phosphorylation in regulating Htt function in health and disease and underscores the critical importance of achieving a more in-depth understanding of the effects of phosphorylation at these residues (31). The knowledge gained from such studies could have important implications for the development of novel therapies and diagnostic strategies based on modulating and/or assessing the levels of phosphorylation at these residues (63). Toward these goals, we utilized protein semisynthetic strategies that were recently developed by our group to generate wild-type and expanded forms of exon1 of the huntingtin protein phosphorylated at S13, S16 or both residues. To allow comparison and correlation of our findings to previously published studies that relied mainly on the use of phosphomimetic mutations, we generated the corresponding phosphomimetic mutant Httex1: S13D, S16D and S13D/S16D. In all cases, we assessed and compared the structure, membrane binding and aggregation properties of each protein to the corresponding unmodified proteins using multiple assays, including sedimentation assays, AFM, TEM and CD spectroscopy. Moreover, we investigated, for the first time, the effect of serine phosphorylation on the internalization and subcellular targeting of Httex1 aggregates in primary striatal cultures.

Previously, biophysical analysis of the effects of N-terminal serine phosphorylation on the *in vitro* aggregation properties of Httex1 utilized peptide model systems in which 40 C-terminal residues of the Httex1 protein were deleted and/or replaced by a solubilizing lysine repeat domain (23,31). Despite these limitations, the results of the aggregation studies using this truncated peptide model are consistent with our observations using tag-free semisynthetic Httex1 proteins and show that serine phosphorylation at either or both S13 and S16 reduce the aggregation rate of the wild-type and mutant Httex1 with diphosphorylation having the strongest inhibitory effect. However, in all previous studies, a direct comparison of the effect phosphorylation and phosphomimetic mutations on the structural properties of Httex1 or Nt17 peptides was never carried out. We postulated that conducting such comparative studies is crucial to determine to what extent the phosphomimetic mutations, which are commonly used to investigate Htt phosphorylation *in vivo*, reproduce all the effects of phosphorylation. Our results demonstrated that phosphorylation at S13 or S16 significantly reduced the rate and extent of wild-type and mutant Httex1 aggregation. Substitution of a single serine by aspartate to mimic phosphorylation failed to reproduce the inhibitory effect of single phosphorylation in the context of mutant Httex1. Previous studies by Wetzel and colleagues only compared the effect of bone fide phosphorylation at S13 and/or S16 to the diphosphomimetics form of mutant Httex1 and showed that the latter exhibits similar aggregation kinetics to the singly phosphorylated proteins (pS13 or pS16) in their Httex1 model system (Nt17-Q37-P10-K2). Our results suggest that single phosphomimetic mutations are not sufficient to reproduce the effect of phosphorylation at single serine residues within Nt17 of mutant Httex1. Phosphorylation of both residues (pS13/pS16) resulted in a marked decrease in the rate of wild-type and mutant Httex1 aggregation and fibril formation. Whereas the S13D/S16D double mutant was able to partially reproduce the inhibitory effect of diphosphorylation pS13/pS16 on the aggregation of mutant **Httex1-Q_42_**, it did not reproduce the effect of phosphorylation on the propensity of **Httex1-Q_42_** to form β-sheet rich aggregate or the effect of phosphorylation on the membrane binding properties of the Nt17 peptide. These results and previous findings by Wetzel and colleagues are consistent with the aggregation inhibitory effects being dominated by charge-mediated mechanisms. The same charge effect was suggested to play a role in Nt17 lipid binding, where it has been shown that acetylation at the 3 lysine residues decreased the binding of Httex1-like peptide model to lipids. However, herein we show that the effects of phosphorylation on membrane binding are strongly dependent on the effect of phosphorylation on the conformation of the Nt17 peptide. Our CD studies showed that phosphorylation at S13 and/or S16 led to complete disruption of Nt17 helix formation in the absence of presence of lipid vesicles and reversed the pT3-induce α-helix formation of Nt17. These findings are consistent with a recent study by Daldin *et al* demonstrating that phosphorylation of S13 and S16 (pS13/pS16-Q_42_) conveys increased conformational flexibility in temperature and polyQ repeat length-dependent experiments (65). They proposed that disruption of the Nt17 α-helix is responsible for this observation. Further studies using NMR and other high-resolution techniques are needed to determine the mechanisms by which phosphorylation-dependent modulation of the conformation of Httex1 contributes to the strong inhibitory effect of this PTM on mutant Httex1 aggregation.

Additionally, serine modifications (pS13, pS16, pS13/pS16 and S13D/S16D) in the context of the Httex1 peptide model were reported to result in the formation of both filamentous and amorphous aggregate species by TEM (23). However, comparable structures were not observed for any of the Httex1 protein in our studies. In contrast, AFM analyses of the aggregates formed mutant pS13, pS16, S13D, S16D, and S13D/S16D Httex1 proteins, showed aggregates of similar morphology to unmodified Httex1 but also revealed differences in the dimensions of these aggregates. It is plausible that these differences in aggregation morphology between the two studies could be caused by the removal of a significant part of the C-terminus, which is known to play important role in enhancing the solubility of Httex1, or by the presence of the non-native lysine residue in the Httex1 model peptide.

A recent study by Branco-Santos suggested that single phosphomimic mutation in mutant Httex1 (97Q) is sufficient to completely abolish Httex1 aggregation and inclusion formation (38). These results are not consistent with our observations and previous studies reported by Wetzel and co-workers (23). The discrepancy between their findings and our results could be due to the differences in the aggregation conditions *(in vitro vs. in cells and in vivo)* or the Httex1 constructs used by both groups. Whereas our studies are based on direct measurements of the aggregation of untagged mutant **Httex1-42Q**, the studies by Branco-Santos *et al* relied primarily on indirect assessment of aggregation using a fluorescence biocomplementation assay (38). Finally, the nature of the aggregates (fibrillar or amorphous) formed by these mutant Httex1-fusion proteins remains unknown,

Next, having generated semisynthetic serine phosphorylated Httex1 proteins, we sought to investigate the effect of *bone fide* phosphorylation on Httex1 aggregate internalization and subcellular targeting in primary striatal neurons. Notably, we found that serine phosphorylation prompted the internalization of aggregated Httex1 and its accumulation within the nuclear compartment. Our results are consistent with a previous study showing that Hek293T cells expressing YFP-Htt-nt17 with phosphomimetic mutations at S13/S16 display enhanced nuclear localization, due to the impaired interaction with the nuclear export protein CRM1 (29). Similar results were observed in the study reported by Thompson et al, which also showed that overexpressed GFP-Htt-ex1-97Q with the S13D-S16D mutations exhibit an increased ratio of nuclear to cytoplasmic fluorescence (6). However, our results demonstrate, for the first time, that genuine phosphorylation of untagged Httex1 fibrils can prompt the accumulation Htt within the nuclear compartment. Moreover, these results suggest that in this case, the phosphomimetics may be reproducing the effect of phosphorylation in directing the subcellular localization of Httex1. These experiments highlight the utility of these protein tools to investigate not only the biophysical consequences of Httex1 phosphorylation but also the interplay between site-selective protein modification, polyQ repeat length, and protein conformation on the biological properties of Httex1 protein, and set the stage for further experimentation with diverse PTMs.

In conclusion, a detailed analysis of the effects of serine phosphorylation on biophysical properties of Httex1 revealed that phosphorylation at serine 13 and/or 16 provokes a remarkable inhibition of aggregation of wild-type and mutant Httex1. A comparative analysis of the effects of the phosphomimetic mutations revealed that the introduction of S→D mutations to mimic phosphorylation at both S13 and S16 significantly reproduces the effect of phosphorylation on Httex1 aggregation, but is insufficient to reproduce the effect of phosphorylation on the conformation of Nt17 and its ability to bind membranes. In primary neurons, the phosphorylated Httex1 proteins appeared to prompt nuclear localization, similar to what has been previously reported with phosphomimetic peptides (6,29). In addition, our studies revealed a novel phosphorylation-dependent molecular switch, involving crosstalk between phosphorylation at T3 and S13 and/or 16, for regulating the conformation of the N-terminus of Htt. Given that phosphorylation at these residues has been implicated in regulating several cellular and toxic properties of Htt, deciphering the PTM-dependent interactome of Htt and unraveling the network of kinases and phosphatases involved in regulating phosphorylation at S13 and S16 should be a top priority to advance our understanding of Htt biology in health and disease. This will facilitate the development of novel strategies to protect against Httex1 aggregation and slow the progression of HD based on selective modulation of Nt17 phosphorylation by regulating Httex1 interactions with other proteins, ligands and cellular compartments.

## Experimental Procedures

### Materials

The peptides used in these studies were synthesized by Integrated Research Biotech Model (IRBM) in collaboration with the CHDI Foundation or purchased from CSBio Co. Glacial acetic acid (AcOH), Trifluoroacetic acid (TFA), guanidine hydrochloride (GuHCl), Tris(2-carboxyethyl)-phosphine hydrochloride (TCEP), methoxyaminehydrochloride (MeONH_4_Cl), 4-mercaptophenylacetic acid (MPAA), 2-methyl-2-propanethiol (*t*-BuSH), sodium borohydride (NaBH_4_), nickel acetate (NiAc_2_), L-proline, L-methionine, and D-trehalose were purchased from Sigma-Aldrich. The 2-2’-Azobis[2-(2-imidazolin-2-yl)propane] dihydrochloride (VA-044) was purchased from Wako. HPLC-grade acetonitrile was purchased from Macherey-Nagel. ER2566 *E.coli* (E6901S) competent cells were purchased from New England Biolabs. Phenylmethanesulfonyl fluoride (PMSF) was purchased from Axonlab. Urea, dimethyl sulfoxide (DMSO), and isopropyl β–D-1-thiogalactopyranoside (IPTG) were purchased from Applichem. CLAP protease inhibitor (1000x) consisted of of 2.5 mg/ml of chymostatin, leupeptin, antipain and pepstatin A from Applichem dissolved in DMSO. The primary mouse anti-Huntingtin monoclonal antibody was purchased from Millipore (MAB5492). Secondary goat anti-mouse antibody labeled with Alexa Fluor 680 was purchased from Invitrogen (A-21057). PageRuler Prestained Protein Ladder (26617) from Thermo Scientific was used for SDS-PAGE. Microcon centrifugal filters with a MWCO of 100 kDa were obtained from Millipore (MRCF0R100). Formvar carbon film on 200-mesh copper grids (FCF200-Cu) and uranyl formate (16984–59-1) from Electron Microscopy Sciences were used for sample preparation for negative-stain transmission electron microscopy (TEM). Liquid chromatography electrospray ionization mass spectrometry (LC-ESI-MS) was performed using a C3 column (Agilent Poroshell 300SB-C3, 1 x 75 mm, 5 µm) with a gradient of 5% to 95% acetonitrile (0.1% v/v formic acid) at 0.3 mL/min over 6 minutes with UV detection at 214 nm and MS detection on a Thermo LTQ. Matrix-assisted laser desorption ionization time-of-flight mass spectrometry (MALDI-TOF-MS) was performed on an AB Sciex 4700 instrument using sinapinic acid matrix. Ultra-performance liquid chromatography (UPLC) analysis was performed on a Waters UPLC system (Waters Acquity H-Class) using a C8 column (Waters Acquity UPLC BEH300, 1.7 μm, 300 Å, 2.1×150 mm) with UV detection at 214 nm using a 2.75-minute gradient of 10% to 90% acetonitrile (0.1% v/v trifluoroacetic acid) at 0.6 mL/min. Preparatory high-performance liquid chromatography (HPLC) was performed on a Waters HPLC system (Waters 2535) using a C4 column (Phenomenex Jupiter C4, 10 μm, 300 Å, 21.2 x 250 mm) with UV detection (Waters 2489) at 214 nm and manual fraction collection.

### Expression and purification of Q18C 18–90 (22Q/42Q) Httex1 protein fragments

Expression of the previously described constructs was performed in *E. coli* BER2566 cells (26). The cells were grown at 37°C until an OD_600_ of 0.1 was reached at which point the incubator was cooled to 14°C. Expression was then induced at this temperature at an OD_600_ of 0.45 with 1 mM (22Q) or 0.4 mM (42Q) IPTG. Bacteria were harvested by centrifugation and lysed by ultrasonication in 22Q lysis buffer (40 mM Tris-acetate, 5 mM EDTA, 30 mM cysteine, pH 8.3, 1x CLAP, and 0.3 mM PMSF) or the 42Q lysis buffer (4.0 M urea, 40 mM Tris-acetate, 5 mM EDTA, 30 mM cysteine, pH 8.3, 1 × CLAP, and 0.3 mM PMSF). The supernatant was separated from the cell debris by centrifugation (23000 × g, 2×20 min, 4°C). For 22Q protein, splicing of the ssp intein was then induced by adjusting the pH to 7.0 and the resulting solution was incubated at room temperature for 3 hours. At this point, or directly after separation from the cell debris for 42Q protein, the supernatant was acidified by the addition of 1% TFA and incubated in boiling water for 5 minutes. The soluble supernatant was separated from the insoluble debris by centrifugation (5000 × g, 15 min, 4°C). After the pH was adjusted to 2.0, the supernatant was filtered (0.2 µm PTFE filter) and purified by preparative HPLC with a gradient from 10 to 40% acetonitrile in H_2_O with 0.1% TFA over 50 minutes after a 10-minute equilibration with 5% acetonitrile. Fractions containing the desired product (elution approximately 40 min) were identified by ESI-MS and analyzed for purity by C8-UPLC. Pure fractions were pooled and frozen in liquid nitrogen and the solvent was removed by lyophilization to yield a white lyophilizate (approximately 4.0 mg/L expression for 22Q protein, approximately 1.5 mg/L expression of 42Q protein).

### Semi-synthesis and desulfurization of wild-type (22Q) Httex1 proteins

Recombinant C-terminal Q18C 18–90 (22Q) was first disaggregated by treatment with neat TFA (~200 µL/mg) for 30 minutes. TFA was removed under a stream of inert gas (N_2_ or Ar) to yield a thin film. The film was then re-suspended by vortexing in ligation buffer solution (6 M GuHCl, 50 mM Tris, 50 mM TCEP, pH 7.0) at ~4 mg/mL§. Methoxyamine was then added (100 mM MeONH_4_Cl), the pH was adjusted to 4.0 and the resulting solution was incubated at 37°C for 1 hour (this treatment was forgone for ligations with peptides containing Ser16 phosphorylation). Next, MPAA (30 mM) was added to the solution and the pH was adjusted to 6.6 or 7.2 (pH 6.6 was used for ligations with peptides containing Ser16 phosphorylation). The corresponding N-terminal NBz peptide fragment (1.5 EQ) was added to the solution, vortexed and the pH was re-checked. The reaction was monitored by LCMS analysis. Upon consumption of the C-terminal Q18C Httex1 18–90 fragment (2–3 hours), the solution was diluted in 6 M GuHCl (to 18 mL), filtered (0.2µm PTFE filter) and purified by preparative HPLC with a gradient from 20 to 55% acetonitrile in H_2_O 0.1% TFA over 50 minutes after a 10 minute equilibration with 5% acetonitrile. Fractions containing the desired products (elution ~35 min) were identified by ESI-MS and analyzed for purity by C8-UPLC. Pure fractions were pooled and frozen in liquid nitrogen, and the solvent was removed by lyophilization to yield a white lyophilizate. The lyophilizate was then disaggregated by treatment with neat TFA (~200 μL/mg) for 30 minutes. TFA was removed under a stream of inert gas (N_2_ or Ar) to yield a thin film. The film was then re-suspended by vortexing in the ligation buffer solution (6 M GuHCl, 50 mM Tris, 500 mM TCEP, pH 7.0) at ~4 mg/mL. VA-044 and t-BuSH were then each added and the resulting solution was incubated at 37°C for 1–2 hours under Ar until LC-MS analysis indicated that desulfurization was completed. The solution was diluted in 6 M GuHCl (to 18 mL), filtered (0.2 µm PTFE) and purified by preparative HPLC with a gradient from 20 to 55% acetonitrile in H_2_O 0.1% TFA over 50 minutes after a 10 minute equilibration at 5% acetonitrile. Fractions containing the desired products (eluted ~35 min) were identified by ESI-MS and analyzed for purity by C8-UPLC. Pure fractions were pooled, frozen in liquid nitrogen, and the solvent was removed by lyophilization to yield a white lyophylizate. (yield of 20–30% over two steps).

### Semi-synthesis and desulfurization of wild-type (42Q) Httex1 proteins

Recombinant C-terminal Q18C 18–90 (42Q) was first disaggregated by treatment with neat TFA (~200 µL/mg) for 30 minutes. TFA was removed under a stream of inert gas (N_2_ or Ar) to yield a thin film. The film was then re-suspended by vortexing in the ligation buffer solution (8 M urea, 500 mM L-proline, 30 mM D-trehalose, 50 mM NaH_2_PO_4_, 50 mM TCEP, pH 7.0) at 1 mg/mL. Methoxyamine was then added (100 mM MeONH_4_Cl), the pH was adjusted to 4.0 and the resulting solution was incubated at 37°C for 1 hour (this treatment was forgone for ligations with peptides containing Ser16 phosphorylation). Then, MPAA (30 mM) was then added to the solution and the pH was adjusted to 6.6 or 7.2 (pH 6.6 was used for ligations with peptides containing Ser16 phosphorylation). The corresponding N-terminal NBz peptide fragment (1.5 EQ) was added to the solution and vortexed and the pH was confirmed and readjusted if necessary to 6.6 or 7.2. The reaction was monitored by LCMS analysis. Upon consumption of the C-terminal Q18C Httex1 18–90 fragment (2–3 hours) the solution was diluted in 8 M Urea (to 18 mL), filtered (0.2 µm PTFE filter) and purified by preparative HPLC with a gradient from 20 to 55% acetonitrile in H_2_O 0.1% TFA over 50 minutes after a 10 minute equilibration with 5% acetonitrile. Fractions containing the desired products (eluted for ~35 min) were identified by ESI-MS and analyzed for purity by C8-UPLC. Pure fractions were pooled and frozen in liquid nitrogen and the solvent was removed by lyophilization to yield a white lyophylizate. The lyophilizate was then disaggregated by treatment with neat TFA (~200 µL/mg) for 30 minutes. TFA was removed under a stream of inert gas (N_2_ or Ar) to yield a thin film. The film was then resuspended by vortexing in the desulfurization buffer solution (20% AcOH, 30 mM L-Methionine, 100 mM TCEP) at 1 mg/mL. Nickel Boride suspension was freshly prepared by reduction of nickel acetate as previously described (50) and mixed with the reaction solution (1:4). The solution was incubated at 37°C with shaking (200 RPM) for 2–3 hours or until LCMS analysis indicated that desulfurization was complete. After The solution was diluted in 6 M GuHCl (to 18 mL), filtered (0.2 µm PTFE filter) and purified by preparative HPLC with a gradient from 20 to 55% acetonitrile in H_2_O 0.1% TFA over 50 minutes after a 10 minute equilibration with 5% acetonitrile. Fractions containing the desired products (eluted ~35 min) were identified by ESI-MS and analyzed for purity by C8-UPLC. Pure fractions were pooled, frozen in liquid nitrogen and the solvent was removed by lyophilization to yield a white lyophilizate. (Yield 15–25% over two-steps).

### In vitro aggregation analysis of Httex1 proteins

Each protein lyophilizate was weighed into an Eppendorf tube and treated with TFA (15 μL/100 µg) for 30 minutes for disaggregation. The TFA was removed under a stream of dry nitrogen. The protein was dissolved in 1X PBS (137 mM NaCl, 2.7 mM KCl, 10 mM Na_2_HPO_4_, 2 mM KH_2_PO_4_) to obtain an approximate concentration of 20 μM and the pH was adjusted to 7.3–4 by the addition of a small volume (1–3 μL) of 1 M NaOH. Any remaining pre-formed aggregates were removed by filtration through 100-kD molecular-weight cutoff filters. The protein concentration was then determined using a UPLC calibration curve based on amino acid analysis (detection at λ_214_). The protein was then diluted to the desired concentration (5 or 3 μM for 22Q and 42Q proteins, respectively) with 1X PBS and confirmed by UPLC analysis and the aggregation experiment was initiated by incubation at the desired temperature (37°C or 25°C for the 22Q and 42Q proteins, respectively). To monitor the soluble protein fraction, the aggregation experiment solution was returned to 25°C and gently mixed. Next, an aliquot (35 μL) was removed at each indicated time-point and aggregates were removed by centrifugation (20,000 × *g*, 20 minutes, 4°C). The supernatant was analyzed by UPLC and the change in the peak area was used to calculate the fraction remaining compared with t=0. Each protein was analyzed in replicate (*n*=4—10) and the combined data from these analyses were plotted and fitted to a single-phase exponential decay. To estimate the rate constant of sedimentation and thus aggregation, we used the function *y* = *Ae*^−*b(x)*^ + *c* where A is the initial concentration of soluble protein, c is the concentration of soluble protein at infinite time and b is the rate constant. We constrained the values of A and c to a range varying between 0 and 100 plus/minus the 2% sensitivity error we commit to the measurement for physical reasons. All fits had an adjusted R^2^ between 0.97–0.99 for **Httex1-Q_22_** curves. All **Httex1-Q_42_** curves had an adjusted R^2^ ranging from 0.93–0.98 (except **S13D/S16D-Q_42_** with R^2^ = 0.90).

### CD analysis

For CD analysis of the aggregation process, aliquots were removed from the aggregation experiments at the indicated time points (100 μL) and analyzed using a Jasco J-815 CD spectrometer in a 1 mm quartz cuvette. The ellipticity was measured from 195 nm – 250 nm at 25°C, and the data points were acquired continuously every 0.2 nm at a speed of 50 nm/min with a digital integration time of 2 s and a bandwidth of 1.0 nm. Five spectra of each sample were obtained and averaged. A sample containing buffer only was analyzed and subtracted from each signal. The obtained spectra were smoothed using a binomial filter with a convolution width of 99 data points and the resulting spectra were plotted as the mean residue molar ellipticity (θ_MRE_).

### Transmission electron microscopy (TEM)

Samples were deposited on Formvar/carbon-coated 200-mesh copper grids (Electron Microscopy Sciences) for 1 min at RT, and then grids were washed and stained with 0.75% w/v uranyl formate (Electron Microscopy Sciences) solution in water. Prepared grids were imaged using a Tecnai Spirit BioTWIN microscope at 80 kV (LaB6 gun, 0.34nm line resolution) equipped with a 4k × 4k Eagle CCD camera with a high-sensitivity scintillator from FEI.

### Atomic force microscopy (AFM) sample preparation, imaging and statistical analysis

Conventional AFM measurements in air of the samples deposited on positively functionalized mica were performed. To functionalize the surface, after cleaving, the bare mica substrate was incubated with a 10 μl drop of 0.05% (v/v) APTES ((3-aminopropyl)triethoxysilane, Fluka) in Milli-Q water for 1 minute at room temperature, rinsed with Milli-Q water and then dried by the passage of a gentle flow of gaseous nitrogen. Preparation of the mica AFM samples was realized at room temperature by the deposition of a 10 µl aliquot of the fully concentrated solution for 10 minutes. Then the sample was then rinsed with ultrapure water and dried with a gentle flow of nitrogen.

High-resolution images (1024×1024 pixels) were collected using an NX10 Atomic Force Microscope (Park Systems, South Korea) under ambient conditions and in amplitude modulation non-contact (NC-AM) mode. We imaged square areas of 2×2 μm2 and 4×4 μm2 areas. We performed all the measurements using ultra-sharp cantilevers (SSS-NCHR, Park Systems, South Korea) with a resonance frequency of 330 kHz and typical apical radius of 2 nm. Raw images were flattened with the built-in software (XEI, Park System, South Korea). To maintain consistency in the further statistical analysis, all images were processed using with the same parameters. First, the images were flattened by a plane and then line by line at a 1st regression order. This second step was repeated until the flat baseline inline profile of the image was reached. During the process of image flattening of the images, the aggregates were masked from the calculation to avoid a modification and underestimation of their height.

Statistical analysis of the cross-sectional dimensions of the fibrillar structures was performed using a home-built program (DNA-trace). In particular, the software enabled tracing of individual fibrils and the measurement of their length and average height.

### Amino acid analysis

Prior to each aggregation experiment 3–5 µg of each protein was dried in an evacuated centrifuge and subjected to amino acid analysis at the Functional Genomic Center Zurich to further confirm that the sample concentration matched the UPLC calibration curve based calculations.

### Membrane binding assay

POPG (1-palmitoyl-2-oleoyl-sn-glycero-3-phospho-(1’-racglycerol), Avanti Polar Lipids) were prepared by extrusion as previously reported (66). The POPG solution was transferred to a glass vial and the solvent was evaporated under a stream of nitrogen. The vial was placed in the lyophilizer under vacuum overnight. The lipid film was resuspended by adding PBS buffer to obtain a multilamellar vesicle suspension with a POPG concentration of 1 mg/mL. The lipids solution was subjected to a water-bath sonication to disrupt the multilamellar vesicles. Large unilamellar vesicles (LUVs) were then prepared by extrusion, applying 21 passes through a polycarbonate filter with 100 nm average pore diameter (Avestin). Samples for membrane binding studies were prepared by adding 5 molar equivalents of lipid to the Nt17 peptides solution (60 μM, final concentration) and incubating the peptide/LUV samples at RT for 1 hour before CD measurements.

### Preparation and characterization of preformed fibrils (PFFs)

To prepare Httex1 PFFs, a 300 µM solution Httex1 was prepared by dissolving lyophilized proteins in 50 mM Tris and 150 mM NaCl, pH 7.5. The resulting solution was centrifuged at 14,000 × *g* (10 min at 4°C) and the supernatant was then incubated at 37°C under constant agitation at 850 rpm on an orbital shaker for 5 days. The generated fibrils were then briefly sonicated (40% amplitude, one pulse for 5 seconds) and then aliquoted and stored at −20°C. Amyloid formation (before and after sonication) was assessed by TEM analysis.

### Primary neuron culture preparation and treatment with preformed fibrils

Primary striatal neuronal cultures were prepared from P0 Sprague Dawley rats as previously described (67). Briefly, dissected tissues were dissociated with papain and triturated using a glass pipette. After centrifugation at 400 × *g* for 2 min, the cells were plated in MEM/10% horse serum onto poly-L-lysine (Sigma) coated coverslips (glass 12 mm, VWR) at 1.5 × 105 cells/ml. The medium was changed after 4 hrs to Neurobasal/ B27 medium. After 7 days in vitro, the neurons were treated with Httex1 PFFs. The concentrations and durations of treatment are indicated in the respective figure legends.

### Western blot analysis of purified Httex1 protein

Protein samples were separated at 180 V on 15% polyacrylamide 1.5 mm-thick gels for 1 h. The proteins were transferred onto nitrocellulose membranes (Omnilab SA) using a semidry transfer system (Bio-Rad) under constant current (500 mA), and the membranes were blocked for 1 h at room temperature with Odyssey blocking buffer (Li-COR Biosciences) diluted 3-fold in PBS buffer. The membranes were probed with the relevant primary antibodies for 4 hours at room temperature. (MAB5492, 1:2000, Millipore; MW8, 1:1000, Developmental Studies Hybridoma Bank). After three washes with PBS buffer containing 0.1% (v/v) Tween 20 (Fluka), the membrane was incubated with secondary goat anti-mouse Alexa^680^ and goat-anti-rabbit Alexa^800^ (dilution 1:2000; Invitrogen). The membranes were then washed three times with PBS buffer containing 0.01% (v/v) Tween 20 and once with PBS buffer, and then they were scanned on a Li-COR scanner at 700 and 800 nm.

### Immunocytochemistry and imaging of primary neurons

As previously described, (78) primary neurons were washed with PBS, fixed with 4% paraformaldehyde in PBS and then incubated in blocking buffer (3% bovine serum albumin (BSA), 0.02% saponin in PBS) for 1 hr at RT. Coverslips were then incubated overnight at 4°C in blocking buffer comprising primary antibodies (MW8, Developmental Studies Hybridoma Bank; MAP2, Cell Signalling). Subsequently, the coverslips were washed five times with PBS and incubated in blocking buffer containing the appropriate Alexa Fluor-conjugated secondary antibodies (Invitrogen) as well as the nuclear stain Hoechst 33342 for 1 hr at RT. After five washes with PBS, the coverslips were washed once with double-distilled water and then mounted on glass slides with DABCO mounting medium (Sigma-Aldrich). The neurons were then imaged using an LSM700 confocal microscope (Carl Zeiss). Image analysis was performed using the Zen 2012 SP1 (black edition, Carl Zeiss) and FIJI softwares.

## Acknowledgment

This work was supported by the CHDI foundation (A_7627) and the Swiss National Science Foundation (31003A-146680). We thank Dr. Sophie Vieweg, Dr. John Warner, Dr. Elizabeth Doherty, Dr. Celia Dominguez and Dr. Andrea Caricasole for helpful discussions on the manuscript, and IRBM for generating the Nt17 peptides used in this study.

The authors declare that they have no conflicts of interest with the contents of this article

